# Early immune pressure imposed by tissue resident memory T cells sculpts tumour evolution in non-small cell lung cancer

**DOI:** 10.1101/2021.04.20.440373

**Authors:** Clare E Weeden, Velimir Gayevskiy, Marie Trussart, Claire Marceaux, Nina Tubau Ribera, Daniel Batey, Charis E Teh, Andrew J Mitchell, Phillip Antippa, Tracy Leong, Daniel Steinfort, Louis Irving, Claire L Gordon, Charles Swanton, Terence P Speed, Daniel HD Gray, Marie-Liesse Asselin-Labat

**Author notes:** Equally contributed to this work (Daniel HD Gray, Marie-Liesse Asselin-Labat).

## Abstract

Tissue-resident memory T cells (T_RM_) provide immune defence against local infection and can inhibit cancer progression. However, it is unclear to what extent chronic inflammation impacts T_RM_ activation and how the immune pressure exerted by T_RM_ affects developing tumours in humans. We performed deep profiling of lung cancers arising in never-smokers (NS) and ever-smokers (ES), finding evidence of enhanced T_RM_ immunosurveillance in ES lung. Only tumours arising in ES patients underwent clonal immune escape, even when evaluating cancers with similar tumour mutational burden to NS patients, suggesting that the timing of immune pressure exerted by T_RM_ is a critical factor in the evolution of tumour immune evasion. Tumours grown in T cell quiescent NS lungs displayed little evidence of immune evasion and had fewer neoantigens with low diversity, paradoxically making them amenable to treatment with agonist of the costimulatory molecule, ICOS. These data demonstrate local environmental insults enhance T_RM_ immunosurveillance of human tissue, shape the evolution of tumour immunogenicity and that this interplay informs effective immunotherapeutic modalities.

## Introduction

Carcinomas evolve from small expansions of transformed epithelial cells to pre-invasive lesions and finally to overt malignancies. All the while, these cells accumulate mutations that can create neoantigens – novel peptides that may provoke an adaptive immune response^1^. The adaptive immune system protects the host from malignant growth by eliminating tumour cells, yet it also exerts selective pressure upon nascent cancers to evade immune predation^2^. Tissue resident memory T cells (T_RM_) provide rapid recall responses to tissue-specific infections and have also been shown to promote cancer-immune equilibrium^3^. How early immunosurveillance by *in situ* tissue-resident T cells might influence immune cell recruitment to developing tumours and impact upon the evolution of tumour immunogenicity is yet to be determined. Lung adenocarcinoma (LUAD) represents an ideal scenario to address these questions due to its prevalence in both never-smoker (NS) and ever-smoker (ES, current- and ex-smoker) patients, where tobacco smoking enhances both the accumulation of DNA alterations in lung epithelial cells^4,5^ and chronic inflammation within the lung^6^. Exploring these two patient groups, we can assess the impact of pre-existing chronic inflammation, tumour mutational burden (TMB) and neoantigen load upon tumour evolution from pre-invasive to invasive late-stage disease.

The genomic drivers of NS LUAD are distinct from those with a history of cigarette smoking. Alterations in *EGFR*, *ROS1* and *ALK* are more common in NS tumours, while *KRAS*, *TP53*, *KEAP1*, *BRAF* and *JAK2/3* alterations are characteristic of ES tumours^7^. These differences guide distinct treatments with targeted kinase inhibitors deployed according to genetic testing^8^. NS patients with lung cancer respond poorly to immune checkpoint inhibitors compared to ES^9^. It has been suggested that tumour cell intrinsic factors such as lower TMB (and presumably lower neoantigen load) and lower expression of PDL1 account for this reduced sensitivity^10–12^. Yet, even in a cohort selected for high expression of PDL1, NS LUAD patients had worse progression free survival and less durable responses to anti-PD(L)1 therapy than heavy smokers^13^. These data imply other smoking-induced factors might also influence checkpoint immunotherapy responses. It was recently found that high densities of tissue-resident memory T cells (T_RM_) within solid tumours were associated with response to anti-PD1 treatment in non-small cell lung cancers (NSCLC), including LUAD and lung squamous cell carcinoma (LUSC)^14^, as well as breast cancers^15^ and melanomas^16^. Therefore we reasoned the presence of T_RM_ or not, and their activation state, may differentially influence tumour evolution in ES and NS patients.

T_RM_ reside in tissues and provide tissue-specific local immunity^17,18^. They have been extensively studied within the murine lung in response to infections such as influenza^19^ and have recently been identified as the dominant T cell population within healthy human lung^20,21^. They can be defined in humans by expression of the memory marker CD45RO, the absence of lymph node homing molecules (i.e. CCR7), surface expression of CD69 and in some instances, the *αε*integrin CD103^22^. An abundance of tumour-infiltrating T_RM_-like cells has been associated with improved prognosis in NSCLC^23,24^. T_RM_ cells are cytotoxic^25^ and secrete pro-inflammatory cytokines, such as IFN-γ and TNF-α, that can recruit and influence the activation of tumour-specific T cells^26,27^. T_RM_ can also activate dendritic cells to increase the numbers of tumour-specific CD8^+^ T cells, conferring protection to tumour cell rechallenge^28^. So-called ‘bystander’ CD8^+^ T_RM_ can therefore support the activity of neoantigen-specific T cells^29^. How T_RM_ within the healthy human lung may be impacted by cigarette smoking and whether these cells play a role in early tumour immunosurveillance is unknown.

Avoiding destruction by the immune system is a hallmark of cancers^30^. ‘Immune escape’ refers to the loss of tumour immunogenicity under T cell predation^31^. Our understanding of the mechanisms of immune escape have focused on tumour intrinsic factors, such as expression of ligands of immune checkpoint receptors (e.g. PDL1), alterations in antigen presentation machinery and neoantigen depletion^32,33^. Loss of heterozygosity (LoH) of the *HLA* locus and neoantigen depletion are observed in untreated NSCLC^34,35^. Neoantigen depletion is also enhanced after treatment with immune checkpoint inhibitors^33^. Recent studies have defined tumour neoantigen evolution as a system dependent on inputs – TMB and immune pressure - influenced by immune escape events to produce neoantigen ‘high’ or ‘low’ tumours^36^. Yet it is unclear whether the pre-cancerous immune environment in which a tumour develops will influence the timing of immune escape events during tumour evolution and has implications for treatment with immunotherapy.

We surveyed non-malignant lung tissue, early-stage primary tumours and late-stage primary tumours in ES and NS patients using mass cytometry and discovered enhanced immunosurveillance by T_RM_ in the lung tissue of ES. This feature translated to greater T cell activation in ES early-stage tumours, even when controlling for TMB. Tumours arising in NS patients did not show evidence of immune pressure until after disease spread to regional lymph nodes. Using multi-region sequencing data from the TRACERx and LxG cohort of early- and late-stage lung cancers^37,38^, we demonstrated clonal immune escape events occurred only in ES tumours, even in those with T cell infiltration and TMB similar to NS patients. These findings suggest that the timing of immune pressure applied by tissue-resident immune cells is a key influence on tumour antigenicity. Finally, we demonstrated that activating co-stimulatory receptors on T cells offers an alternative immunotherapeutic strategy for T cell quiescent tumours such as those occurring in NS patients.

### Mass cytometric profiling of non-malignant lung and early-stage NSCLC reveals distinct immune landscapes in never smoker and smoker patients

We first examined the global immune cell composition in early-stage NSCLC tumours and adjacent non-malignant lung tissue from 20 patients (10 NS, 10 ES, Supp Table 1) by mass cytometry (CyTOF) and immunohistology. A purpose-built panel of 43 CyTOF antibodies was employed for deep profiling of tumour, haematopoietic and tissue resident lymphoid cells (Supp Table 2). All antibodies were validated to ensure specificity (e.g. Supp Fig 1a) and key determinants were correlated with gold standard immunohistochemical detection (Supp Fig 1b,c). The percentage of CD45^+^ cell infiltration was consistent within non-malignant tissue but more variable in tumours, with some tumours having a significantly lower immune infiltration than matched normal tissue (Supp Fig 1b). Data were corrected for batch effects^39^ and the CD45^+^ infiltrate classified using FlowSOM clustering^40^ to identify 14 cell populations encompassing lymphoid and myeloid cell lineages and visualized using tSNE dimensionality reduction (Fig 1a, Supp Fig 1d). Each cluster was represented to varying degrees in non-malignant and tumour tissue, in both ES and NS and in each individual patient (Fig 1b,c and Supp Fig 1e). When comparing tumours and corresponding non-malignant tissue, high inter-patient variability was apparent and significant changes in the proportions of CD4^+^ T cells, NK cells, macrophages and neutrophils were detected (Supp Fig 1e).

**Figure 1.**
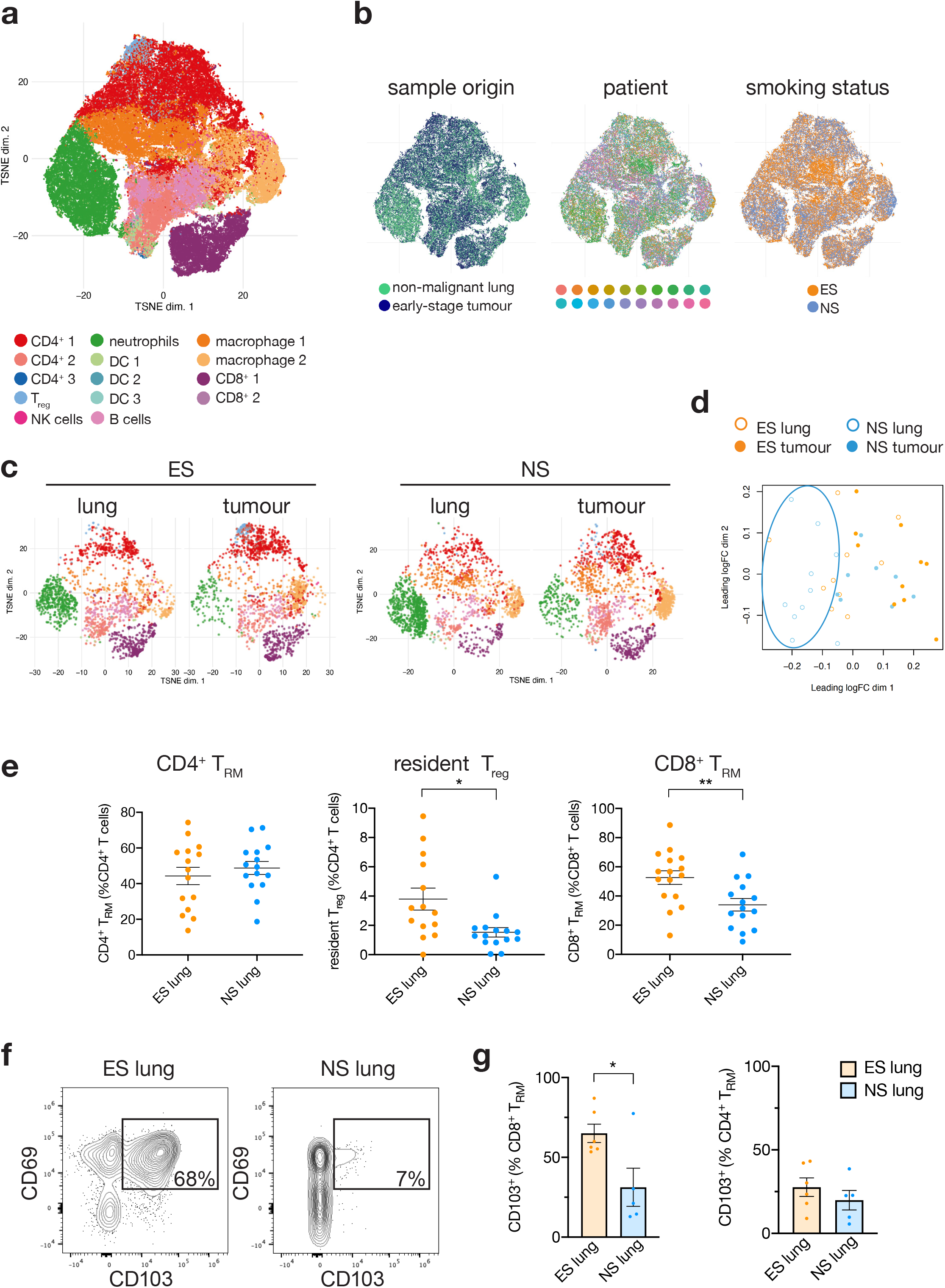
The immune micro-environment in non-malignant lung tissue and early-stage lung cancers. **a**, FlowSOM clustering and tSNE visualisation of CD45^+^ cells from 41 samples (20 patients, total 82,000 cells total). **b**, tSNE visualisation of CD45^+^ cells profiled by CyTOF from non-malignant lung tissue and early-stage lung cancer, coloured by sample origin, patient, and smoking status as indicated. **c**, Representative tSNEs of CD45^+^ cells from an ever-smoker patient (MH0136, 47 yo female, current smoker, stage I LUAD) and a never-smoker patient (SH0139, 68 yo female, stage I LUAD). **d**, Multi-dimensional scaling plot of Boolean gated data of CD45^+^ cell phenotypes in non-malignant lung and tumour from ES (orange) and NS (blue) patients. Manually defined ellipse highlights non-malignant lung (blue line) cluster in NS patients. **e,** Proportion of CD3^+^CD69^+^CD45RO^+^CCR7^-^ T_RM_ cells in CD4^+^ T cells, T_reg_ and CD8^+^ T cells in non-malignant lung tissue analysed by CyTOF and flow cytometry. n = 16 ES, n = 15 NS. Unpaired t-test, *p<0.05, **p<0.01. **f,** Representative flow cytometry plots showing CD103 versus CD69 expression, gated on CD3^+^CD45RO^+^CCR7^-^CD8^+^ T_EM_ from non-malignant lung tissue. ES patient (MH0070, 56 yo female, current smoker, LUAD) and NS patient (MH0066, 77yo male, LUAD). **g**, Quantification of the proportion of CD103^+^ cells within the CD3^+^CD69^+^CD45RO^+^CCR7^-^ CD8^+^ T_RM_ or CD4^+^ T_RM_ compartment by flow cytometric analysis of non-malignant lung tissue, n= 6 ES, n=5 NS, unpaired t-test, *p<0.05.

Fine mapping of the immune landscape was performed using conventional gating of immune cell populations with phenotyping markers, which were then expressed as a proportion of each parent immune population. This approach generated 234 immune parameters per patient (Supp Table 2) and we produced multi-dimensional scaling plots of this data using the R package limma^41^. We found that the immune cell features from non-malignant lung tissue in NS patients were largely segregated from all the other samples (Fig 1d). In contrast, CD45^+^ haematopoietic cells in normal ES lung were more closely correlated to tumour samples (Fig 1d), implying that smoking dramatically alters immune phenotypes within the lung. These results prompted us to compare the resident T cell populations in non-malignant ES and NS lung tissue. We found no significant difference in the overall proportion of CD4^+^ T_RM_ cells (CD69^+^CD45RO^+^CCR7^-^ cells, as defined in Supp Table 4) between ES and NS non-malignant lungs in 31 patients analysed by CyTOF or flow cytometry (Fig 1e and Supp Table 1,2,3; n = 16 ES and n = 15 NS). However, we observed a striking increase in the frequencies of CD8^+^ T_RM_ and resident T_reg_ cells in ES lung tissue (Fig 1e). Human lung T_RM_ cells can be further subdivided by CD103 expression, where CD103^+^ cells express higher expression of activation markers compared to CD103^-^ cells^21,42^. We found more than double the proportion of CD103^+^CD69^+^ T cells within the CD8^+^ T_RM_ cell population in ES lung compared to NS patients (Fig 1f,g, and Supp Table 4). Importantly, the proportions of T_RM_ we found in the lung tissue of cancer patients were comparable to those observed in healthy lung of non-cancer patients^43^, indicating analysis of non-malignant lung tissue reflects the state of the immune microenvironment prior to tumour formation. Overall, these results demonstrate that the prevalence of resident T_reg_ and CD8^+^ T_RM_ distinguishes ES non-malignant lungs from NS tissue.

### Tissue resident memory T cell immunosurveillance is diminished in non-malignant lung tissue of never-smoker patients

Engagement of T cell accessory molecules determines the functional outcome of TCR signalling. Their expression can be used to infer T cell activation state and tissue-specific immunosurveillance^44^. In the early phase of T cell activation, expression of co-stimulatory molecules such as CD28, CD27, 41BB and OX40 predominates and their engagement can amplify T cell responses to antigens. At the peak of the T cell response, both co-stimulatory and co-inhibitory molecules are expressed followed by a preponderance of co-inhibitory receptors (CD57, PD1, LAG3, TIM3) towards the end of the response^44^. Global analysis of the expression of accessory molecules in lung resident T cells revealed an abundance of co-stimulatory molecules in ES non-malignant lung that was not observed in NS lungs (Fig 2a). Specifically, enhanced expression of CD27, OX40, CTLA4 and ICOS was detected on CD4^+^ T_RM_ in ES patients (Fig 2b,c). In ES lung tissue, more CD8^+^ T_RM_ expressed Ki67 and perforin, indicating increased proliferation and cytotoxicity of these cells in ES lungs compared to NS tissue (Fig 2d), although no consistent differences were detected in the expression of accessory molecules in resident T_reg_ and CD8^+^ T_RM_ cells between ES and NS tissue (Supp Fig 2a). To further evaluate the activation status of T cells in ES and NS lungs, T cells purified from non-malignant tissue were stimulated with anti-CD3 and anti-CD28 and assayed for proliferation 48h and 72h after TCR engagement. CD4^+^ and CD8^+^ T cells isolated from NS lung displayed markedly reduced proliferation compared with ES CD4^+^ and CD8^+^ T-cells (Fig 2e,f). Taken together, these data provide compelling evidence that there is greater T_RM_ cell immunosurveillance within ES lungs and lung T cells are poised for effector function in ES patients, but not in NS lungs.

**Figure 2.**
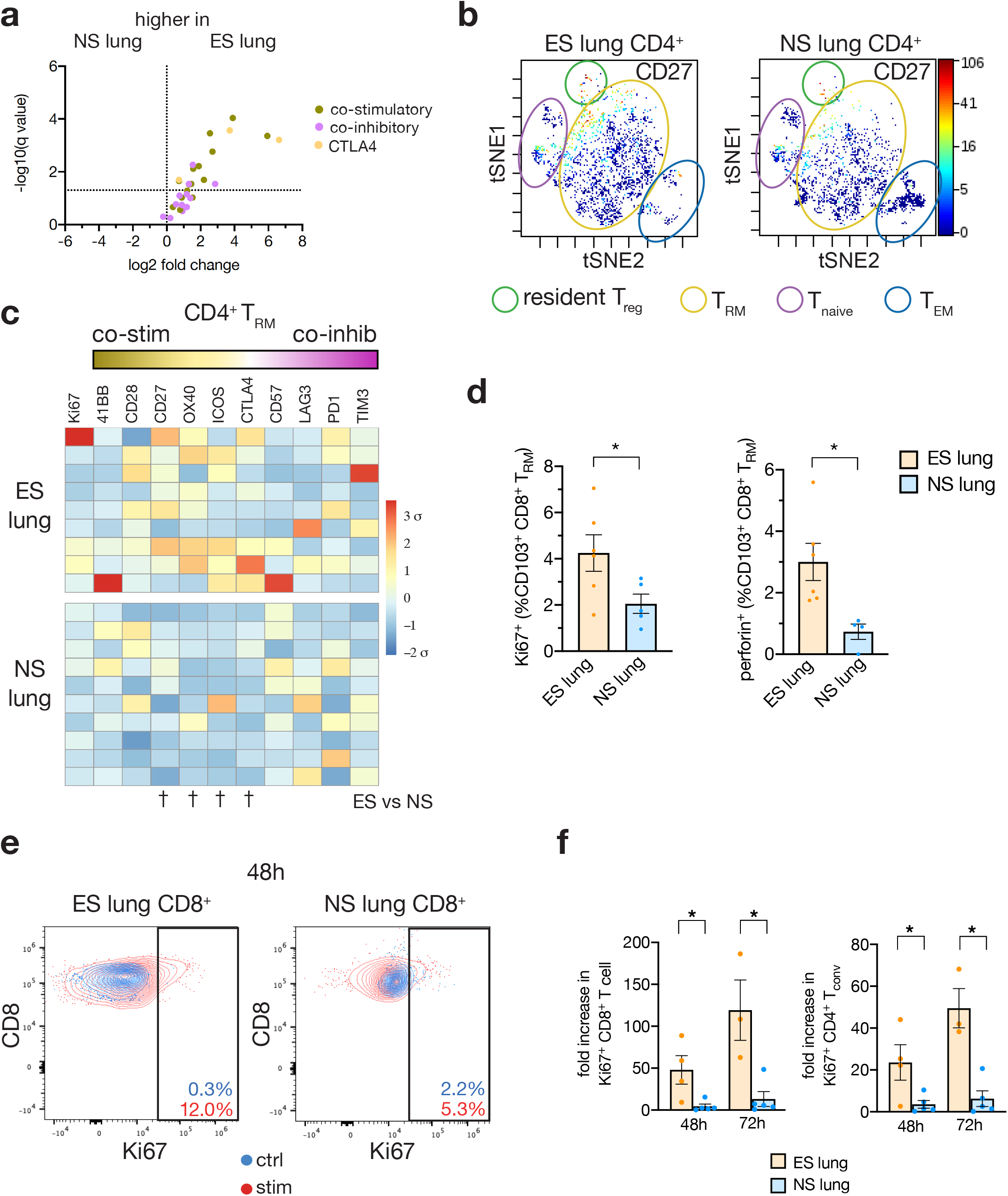
Lung resident memory T cells are less activated in never-smoker lung tissue. **a**, Volcano plot depicting fold change in T_RM_ accessory molecules between NS and ES lung. 30 parameters analysed by CyTOF, co-stimulatory molecules in green, CTLA4 in yellow and co-inhibitory markers in purple. Dotted line indicates q value = 0.05. Data represents average marker fold change within each cell type (CD4^+^ T_RM_, resident Treg and CD8^+^ T_RM_), n = 10 ES and n = 8 NS patients. **b,** Representative viSNE analysis of CD4^+^ T cells in non-malignant lung tissue showing CD27 expression overlayed in an ES patient (MH0103, 58 yo male, ex-smoker, stage IIIa LUAD) and NS patient (AH0059, 79yo female, stage I LUAD). **c**, Heatmap of co-stimulatory and co-inhibitory molecule expression on CD69^+^CD45RO^+^CCR7^-^CD4^+^ T_RM_ in non-malignant lung tissue analysed by CyTOF. Data is scaled to mean expression ± standard deviation per marker. N = 9 ES and n = 10 NS patients. Unpaired t-test, †p<0.05. **d**, Quantification of the proportion of Ki67^+^ and perforin^+^ cells within the CD3^+^CD69^+^CD103^+^CD8^+^ T_RM_ compartment by flow cytometric analysis of non-malignant lung tissue, n= 6 ES, n=5 NS, unpaired t-test, *p<0.05. **e**, Representative flow cytometry plots of CD8 versus Ki67 expression 48h after anti-CD3/anti-CD28 stimulation of CD8^+^ T cells sort purified from non-malignant lung tissue. ES patient (MH0067, 47 yo female, ex-smoker, LUAD) and NS patient (MH0044, 47 yo male, carcinoid). **f**, Fold change in the proportion of Ki67^+^ CD4^+^ and CD8^+^ T cells at 48h and 72h after anti-CD3/anti-CD28 stimulation of T cells sorted from non-malignant lung tissue. n = 4 ES; n = 5 NS; unpaired t-test, *p<0.05.

### CD8+ T cells have limited activation in early stage never-smoker lung adenocarcinoma

The markedly low activation potential of resident T cells in NS lungs led us to assess their phenotype during tumourigenesis. CD103 and CD69 expression are not reliable correlates of tissue residency within tumours as these markers are transiently upregulated in response to inflammation and TGFß, yet they can still be used to identify activated T cells in this setting^45,46^. The proportion of CD4^+^ T_RM_-like cells was not consistently altered among tumour-infiltrating T cells, yet enrichment of CD69^+^ T_reg_ cells was observed within tumours compared with non-malignant lung, irrespective of smoking status, consistent with previous studies^47^ (Supp Fig 3a,b,c). Increased proportions of CD8^+^ T_RM_-like cells were found in tumour tissue from NS patients but not from ES patients, likely due to the already enhanced CD8^+^ T_RM_ presence in ES lungs (Supp Fig 3d).

Phenotypic analysis of these T cells revealed marked upregulation of accessory molecules in CD4^+^, T_reg_ and CD8^+^ T_RM_-like cells infiltrating ES tumours compared to matched lung tissue (Fig 3a). In contrast, only a few instances of modest increases in co-stimulatory molecule expression were observed in CD4^+^ and T_reg_ T_RM_-like cells in NS tumours compared to matched non-malignant lung (Fig 3a). The expression of co-inhibitory molecules remained low in both tissues from NS patients (Fig 3a). Moreover, the expression of accessory molecules in CD8^+^ T_RM_-like cells infiltrating NS tumours remained low, similar to the level observed in non-malignant NS lungs (Fig 3a). When comparing the range of T cell phenotypes analysed, enrichment in cells expressing co-stimulatory and co-inhibitory receptors was exclusively found in ES tumours compared with NS tumours (Fig 3b). Further interrogation of CD8^+^ T_RM_-like phenotypes revealed features of exhausted T cells^48^ (defined by co-expression of multiple co-inhibitory markers including PD1, TIM3 and LAG3) in ES tumours, features that were not detected in NS tumours (Fig 3c,d). We next addressed whether the low activation status of CD8^+^ T cells found in our cohort of NS patient tumours was detectable in a larger cohort. We found evidence of lower CD8^+^ T cells infiltration of NS tumours using three different algorithms to detect the transcriptional signatures of CD8^+^ T cell infiltration and activation within bulk RNAseq data from TCGA LUAD (Fig 3e). These data establish that T cells infiltrating NS tumours are largely quiescent.

**Figure 3.**
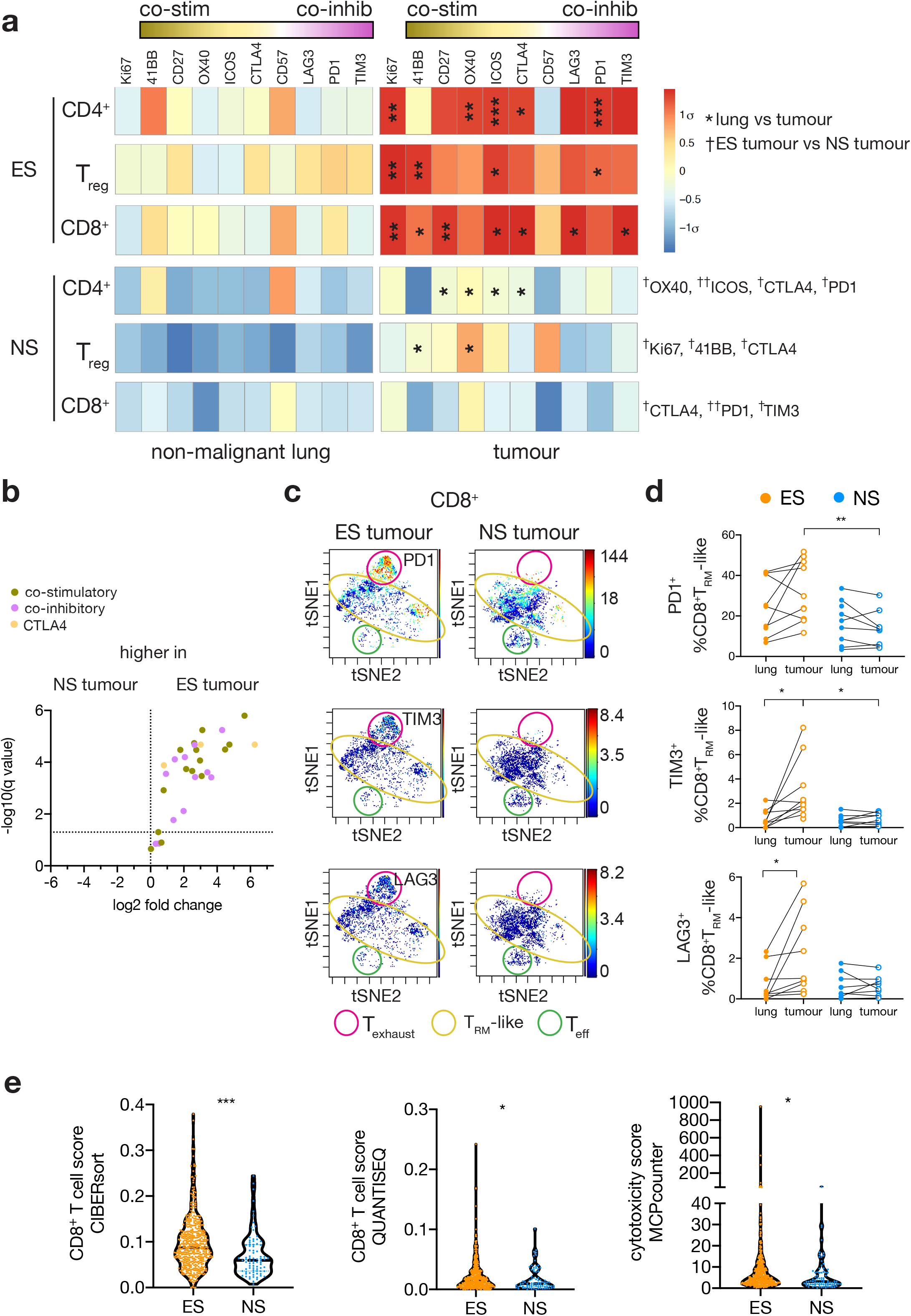
Tumour-associated T cells have limited activation in early-stage never-smoker lung cancer. **a**, Heatmap of the mean proportion of CD3^+^CD69^+^CD45RO^+^CCR7^-^ CD4^+^, T_reg_ and CD8^+^ T cells expressing co-stimulatory and co-inhibitory proteins as detected by CyTOF. Data is scaled to mean expression ± standard deviation per marker. n = 10 ES non-malignant lung, n = 10 ES tumours; n = 8 NS non-malignant lung, n = 8 NS tumours. Paired t-test between lung and tumour, *p<0.05, **p<0.01, ***p<0.001. Unpaired t-test between NS and ES tumours, †p<0.05, ††p<0.01. **b**, Volcano plot depicting fold change in T_RM_ accessory molecules between NS and ES tumours. 30 parameters analysed by CyTOF, co-stimulatory molecules in green, CTLA4 in yellow and co-inhibitory markers in purple. Dotted line indicates q value = 0.05. Data represents average marker fold change within each cell type (CD4^+^ T_RM_-like, resident-like Treg, and CD8^+^ T_RM_-like), n = 10 ES and n = 8 NS tumours. **c**, Representative viSNE analysis of CD8^+^ T cells in tumour tissue, overlayed with PD1 (top panel), TIM3 (middle panel) or LAG3 expression (bottom panel). ES patient (MH0136, 47yo female, current smoker, stage I LUAD) and NS patient (AH0089, 60 yo female, Stage IIIa LUAD). **d**, Quantification of the proportion of PD1^+^, TIM3^+^, or LAG3^+^ cells among CD3^+^CD69^+^CD45RO^+^CCR7^-^ CD8^+^ T_RM_-like cells as detected by CyTOF. n = 10 ES, n = 8 NS. Paired t-test between lung and tumour, unpaired t-test between NS and ES. *p<0.05, **p<0.01. **e**, Violin plots depicting CD8^+^ T cell scores detected by three different algorithms from bulk RNAseq of The Cancer Genome Atlas LUAD cohort using TIMER2.0^85^. n=472 ES, n=83 NS. Unpaired t-test with Welch’s correction *p<0.05, ***p<0.001.

### Delayed and reduced immune pressure in NS tumours compared to ES tumours of similar mutational burden

Our analyses revealed a significant enhancement in T_RM_ immunosurveillance in the lungs of ES, correlating with greater T cell activation in early-stage tumours. A major determinant of immune cell recruitment to cancers is tumour mutational burden (TMB)^49^. Cancer cells with a higher TMB produce more neoantigens that have the potential to be presented and recognised by T cells. Therefore, we examined whether the enhanced T cell activation we detected in early-stage ES tumours was solely attributable to greater TMB in these patients compared to NS^7,10,11^. TMB positively correlated with the predicted number of neoantigens (Supp Fig 4a) and both values were low in NS patients compared with ES patients in the TCGA LUAD^50^ and TRACERx cohort of LUAD and NSCLC not otherwise specified (NSCLC-NOS, detailed in^37^, Supp Fig 4b-e). We also detected positive associations between both TMB and the number of predicted neoantigens with CD8^+^ T cell score in the TCGA LUAD dataset (Supp Fig 4f,g), demonstrating the importance of controlling for this variable. When comparing NS and ES tumours with similar TMB (< 4.5 mt/Mb, Fig 4a,c), NS tumours still had lower immune recruitment scores than ES TMB^lo^ tumours in these two datasets (Fig 4b,d). In addition, CD8^+^ T cell scores were also lower in NS pre-invasive adenocarcinoma lesions than ES, where TMB are below 4.5 mt/Mb in both groups (mean TMB for ES = 1 mt/Mb; NS = 0.8 mt/Mb, Fig 4e) ^51^. These data suggest that immune cell recruitment in pre-invasive and early-stage LUAD is not only associated with TMB but also with patient smoking status.

**Figure 4.**
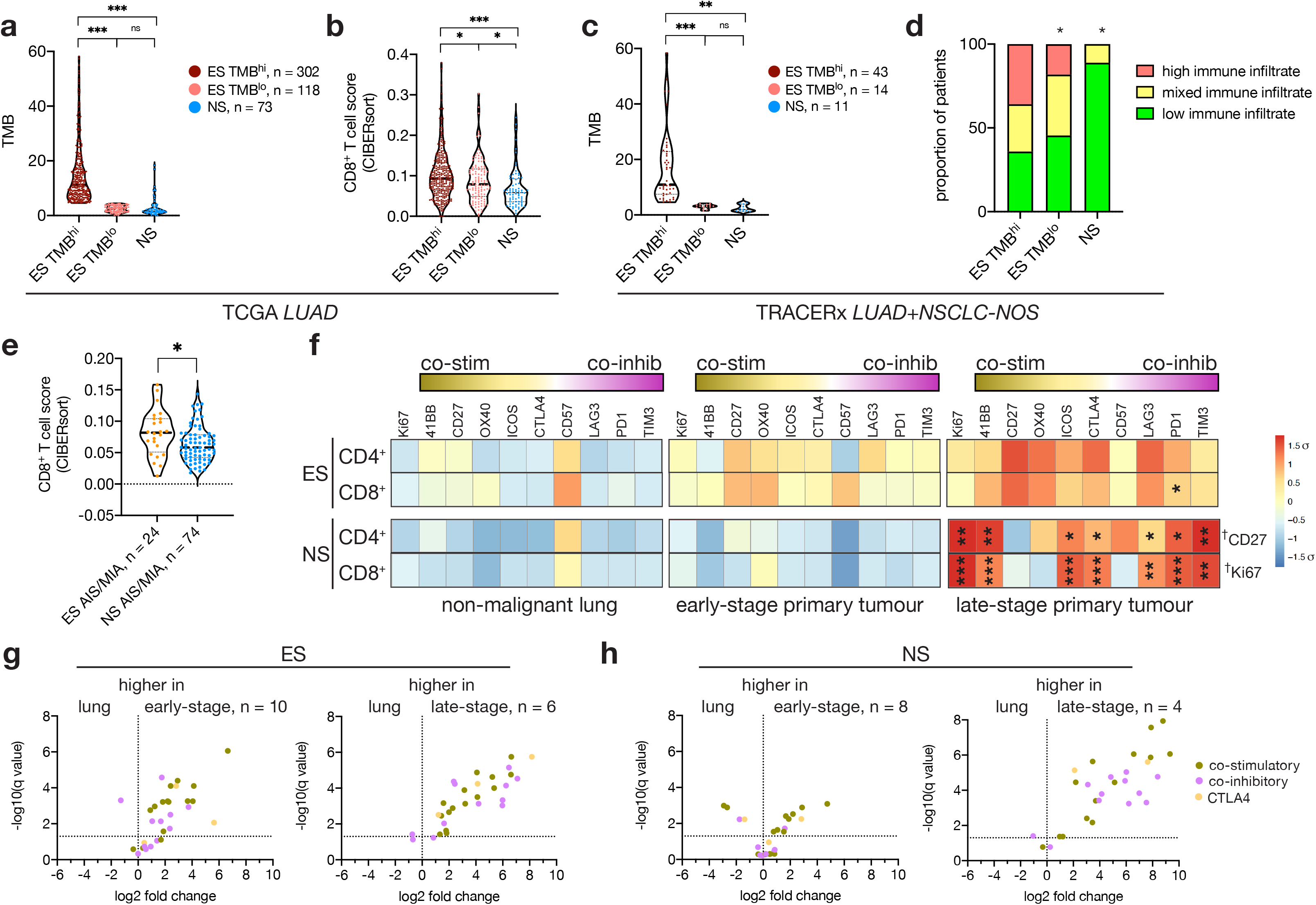
Smoking enhances early immune recruitment to tumours independent of tumour mutational burden. **a**, Measure of tumour mutational burden (TMB) from whole exome sequencing data from the TCGA LUAD cohort related to smoking status, as calculated in Hoadley *et al.*^84^. Unpaired t-test with Welch’s correction. *p<0.05, ***p<0.001. **b**, CD8^+^ T cell scores estimated using RNA sequencing data from the TCGA LUAD cohort as in Fig 3e related to patient smoking. **c**, Average TMB per patient (calculated as the TMB summed for all sites/# samples) in the TRACERx cohort of LUAD and NSCLC not otherwise specified (NSCLC-NOS). Unpaired t-test with Welch’s correction. **p<0.01, ***p<0.001. **d**, Immune cell scores estimated using Danaher signatures extracted from multi-site RNA sequencing from TRACERx patients as detailed in Rosenthal *et al.*^37^. Fisher’s exact test, *p<0.05. **e,** CD8^+^ T cells scores estimated from bulk RNAseq data from the Chen cohort of lung adenocarcinoma *in situ* (AIS) and minimally invasive adenocarcinoma (MIA) ^90^. Unpaired t-test with Welch’s correction. *p<0.05. **f,** Heatmap of the mean proportion of co-stimulatory and co-inhibitory protein expressing cells within CD3^+^CD69^+^CD45RO^+^CCR7^-^ CD4^+^ T_RM_-like, resident-like Treg, and CD8^+^ T_RM_-like cells detected by CyTOF. Data is scaled to mean expression ± standard deviation per marker. n = 10 ES non-malignant lung, n = 10 ES stage I-IIIa primary tumours (early-stage); n = 6 ES stage III/IV primary tumours (late-stage); n = 10 NS non-malignant lung, n = 8 NS stage I-IIIa tumours (early-stage); n = 5 NS stage III/IV primary tumours (late-stage). Matched non-malignant lung and stage I-IIIa tumours acquired from surgically resected tissue; stage IIIb/IV tumours acquired from endobronchial ultrasound guided biopsies of primary tumours from an independent cohort of patients. Unpaired t-test between early-stage and late-stage disease, *p<0.05, **p<0.01, ***p<0.001. Unpaired t-test between late-stage tumours from ES and NS patients, †p<0.05. **g,h,** Volcano plot depicting fold change in T_RM_ accessory molecules between lung and tumours. 30 parameters analysed by CyTOF, co-stimulatory molecules in green, CTLA4 in yellow and co-inhibitory markers in purple. Dotted line indicates q value = 0.05. Data represents average marker fold change within each cell type (CD4^+^ T_RM_-like, resident-like Treg, and CD8^+^ T_RM_-like), in ES (**g**) and NS (**h**) samples. n values as indicated, n = 10 lung samples for both ES and NS.

We next sought to address whether smoking-induced differences in T cell activation were maintained throughout tumour evolution. We assessed T cell phenotypes in late-stage biopsies of primary lung cancers from a further 7 ES and 5 NS patients, using CyTOF to correlate immune pressure with disease stage (n = 10 ES Stage I-IIIa, n = 10 NS Stage I-IIIa; n = 7 ES Stage III/IV and n = 5 NS Stage III/IV, Supp Table 1).). In ES patients, both CD4^+^ and CD8^+^ T cells exhibited a gradient of increased activation from non-malignant tissue to early-stage then to late-stage cancers (Fig 4f). However, T cells within NS patients did not show evidence of activation until late-stage disease (Fig 4f), by which point smoking-induced differences in the expression of T cell accessory molecules were less apparent (Fig 4f, Supp Fig 4h). This pattern was also evident when plotting the overall changes in co-stimulatory and co-inhibitory molecules. In ES patients, almost all increased in both early- and late-stage tumours (Fig 4g), whereas in NS patients, they were only substantially higher in late-stage cancers (Fig 4h). This activation profile suggests that tumours arising in ES not only experience enhanced immune pressure, but that this occurs earlier than in NS patients - perhaps related to the early presence of activated T_RM_ cells in ES non-malignant lung tissue.

### Delayed immune pressure in NS patients results in minimal immune escape of tumour clones

Recent studies have defined the system of tumour neoantigen evolution as one dependent upon inputs, including (i) TMB and (ii) the level of immune selective pressure, that influence immune escape events such as (iii) loss of heterozygosity of *HLA-A*,-*B* or -*C* alleles (*HLA* LoH) and (iv) depletion of neoantigens or immunoediting^36^. The system results in the output of (v) clonal neoantigens – neoantigens that are most likely to elicit T cell activity and are associated with successful responses to checkpoint immunotherapy^52^. Given the substantial differences in TMB and immune pressure identified with smoking status, lung cancer occurring in ES and NS patients provides an opportunity to define the evolution of tumour immunogenicity using real world data. To infer the timing of immune escape events such as *HLA* LoH and immunoediting in NS and ES tumours, we used multi-site whole exome sequencing data from the TRACERx cohort of early-stage, surgically resected LUAD, LUSC and NSCLC-NOS tumours^37^ and whole genome sequencing data from the LxG cohort of late-stage, biopsied LUAD, LUSC and SCLC tumours and metastases^38^. Clonal or truncal events (present in all sites) occur earlier in tumour evolution compared to those described as subclonal that are shared (present in some sites) or private (present in only one site).

In both datasets, clonal *HLA* LoH was detected only in ES patients, irrespective of cancer type or stage (Fig 5a and Supp Fig 5a). Intriguingly, clonal *HLA* LoH was also apparent in tumours from ES patients with low TMB and with low immune infiltrate (Supp Fig 5b,c), yet not detected in a late-stage NS patient, who likely had high T cell activation (Fig 5a, Fig 4e). Moreover, analysis of *HLA* LoH in an independent cohort of pre-cancerous forms of LUAD, adenocarcinoma *in situ* and minimally invasive adenocarcinoma (AIS/MIA^51^) also confirmed that this event was rare in NS patients (Supp Fig 5d). These data show that immune pressure present from the initiation of tumourogenesis in ES lungs can precipitate early tumour immune escape events, suggesting the timing of immune pressure influences tumour immune evasion in addition to the level of immune activation.

**Figure 5.**
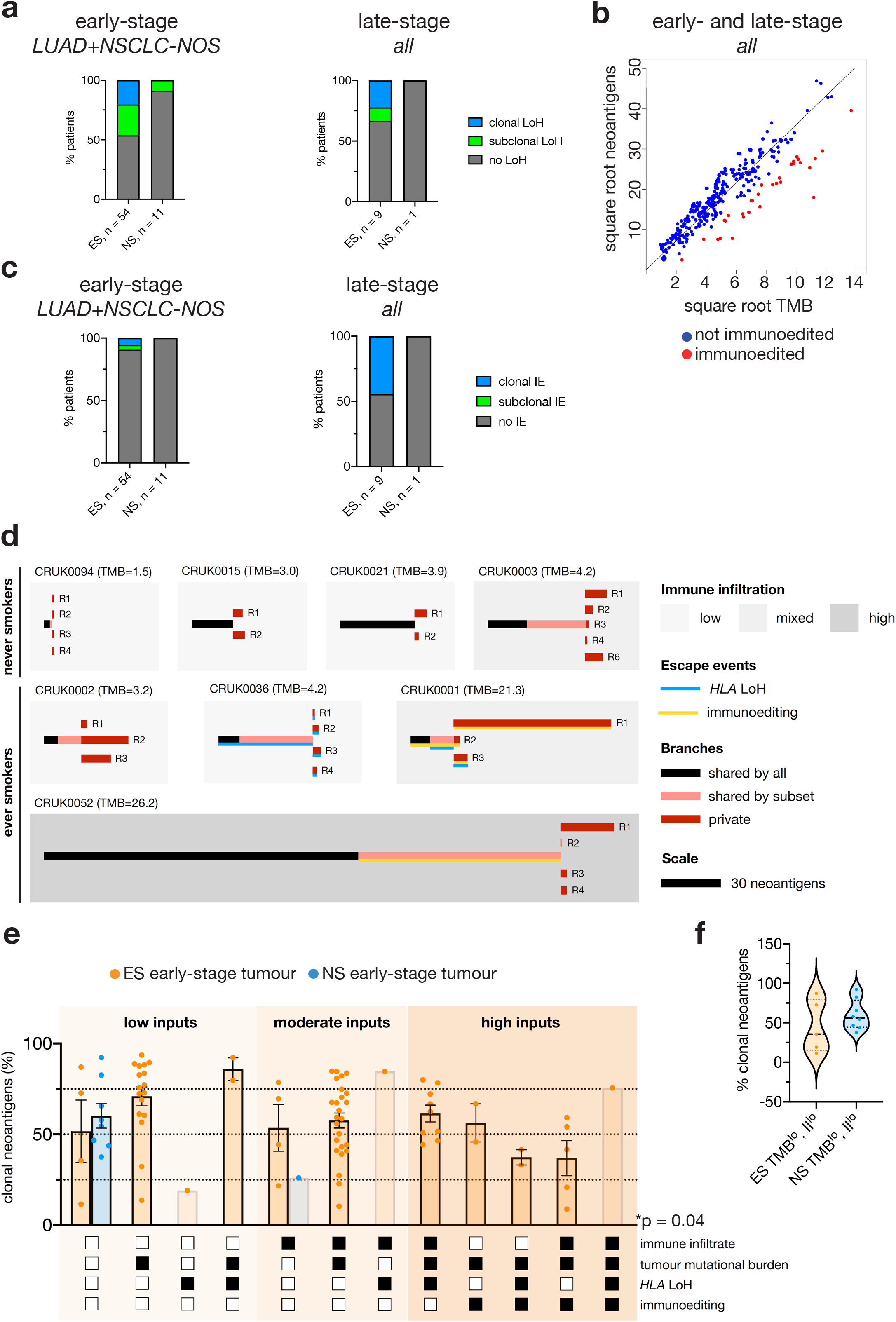
The timing of immune pressure influences tumour neoantigen evolution in lung cancer. **a**, Proportion of patients with loss of heterozygosity of one or more *HLA* alleles in some tumour sites (subclonal LoH; green) or all sites (clonal LoH; blue) in the TRACERx cohort of LUAD and NSCLC-NOS (early-stage, left panel) and the LxG cohort of metastatic LUAD, LUSC and SCLC (late-stage, right panel). Fisher’s exact test, p = 0.07 of TRACERx cohort. Fisher’s exact test not possible for LxG cohort due to only 1 NS patient. **b**, Scatter plot showing the power transformed square root of tumour mutational burden and neoantigen numbers from samples within the TRACERx and LxG cohorts. The trend line is fitted using robust linear modelling. Samples with weights less than 0.9 and negative residuals are called as ‘immunoedited’ and indicated in red, detailed in Methods. **c**, Proportion of patients with fewer than expected numbers of neoantigens for a given tumour mutational burden in the TRACERx (early-stage, left panel) and LxG cohort (late-stage, right panel). Immunoediting (IE) present in all sites is termed clonal IE while that observed in only some tumour sites is termed subclonal IE. Fisher’s exact test, not significant. **d**, Neoantigen trees depicting neoantigens shared by all sites (black), present in some sites (pink) or present in only one site (red) in representative ES and NS patients in the TRACERx cohort. The length of the trunk and branches is scaled to the number of neoantigens. Background colour indicates the level of immune infiltration as determined from RNA sequencing. LoH of *HLA-A*, -*B* or -*C* is indicated in blue, immunoediting is indicated in yellow, and the position of the line represents clonal or subclonal occurrence. **e**, Proportion of clonal neoantigens in the TRACERx cohort of ES (orange) and NS patients (blue). Patients are separated according to tumour inputs – hi/mixed immune infiltrate and high tumour mutational burden indicated by filled black box. The presence of clonal *HLA* LoH or clonal/subclonal immunoediting are indicated by filled black box. Kruskal-Wallis test, *p<0.05. **f**, Violin plot depicting the range of clonal neoantigens within TRACERx patients harbouring low immune infiltration and low tumour mutational burdens. Levene’s test of variance, p=0.07.

We next defined ‘immunoedited’ or neoantigen depleted samples as those TRACERx and LxG samples with fewer than expected neoantigens for a given TMB using robust linear modelling (see Methods, Fig 5b). Samples with low weights (i.e. distant from the consensus line) were more likely to have negative residuals (Supp Fig 5e). This finding demonstrates that amongst outliers, our dataset consists of largely neoantigen-depleted samples rather than neoantigen-enriched samples; further evidence of immunoediting. Immunoediting was uncommon, yet clonal neoantigen depletion was only found in patients with high/mixed immune infiltrate or high TMB and was unrelated to *HLA* LoH events (Supp Fig 5f). Consistent with the *HLA* LoH data, clonal immunoediting was only detected in ES patients (Fig 5c, Supp Fig 5f), providing further evidence of early immune recognition and depletion of immunogenic tumour clones in ES compared to NS patients.

Finally, we assessed how each system input – TMB and immune infiltrate – and immune escape events affected the proportion of clonal neoantigens detected in early-stage lung cancers. Only tumours with immunoediting were associated with significantly reduced proportions of clonal neoantigens (Supp Fig 5g). No significant differences were detected in relation to TMB, immune infiltrate nor *HLA* LoH events (Supp Fig 5g). Overall, all tumours occurring in the NS patient cohort had low TMB and immune infiltrate, resulting in rare subclonal immune escape events and thus relatively high proportions of clonal neoantigens in the majority of patients (Fig 5d,e). Only one NS patient, CRUK0003, with mixed immune infiltrate had clonal neoantigens close to 25%, and all other NS patients had simple tumour-immune evolutionary trees (Fig 5d,e). In contrast, a full gamut of TMB and immune infiltrates in ES tumours resulted in frequent clonal and subclonal immune escape events and thus divergent tumour-immune pedigrees within the ES cohort (Fig 5d,e). Indeed, classifying combinations of system inputs and immune escape resulted in 12 distinct categories detected within the ES cohort compared to only 2 in NS patients (Fig 5e). Higher TMB and immune pressure was more likely to lead to immune escape in these ES patients, and thus proportionally fewer clonal neoantigens (Fig 5d *eg*. CRUK0001 and CRUK0052; Fig 5e, ‘high inputs’). Critically, even among ES patients with TMB and immune infiltrates similar to NS patients, we found ES patients with immune escape events, resulting in a broad range of clonal neoantigen proportions (Fig 5d *eg*. CRUK0002, CRUK0036, Fig 5f). These data suggest that the timing of immune pressure is an important determinant of tumour immunogenicity in addition to level of immune infiltration and TMB.

The finding that delayed immune pressure in NS patients leads to less immune escape and higher frequency of clonal neoantigens, albeit at lower numbers, suggests that these patients could potentially engage T cell responses targeting most tumour clones. However, promoting such responses would require an alternative strategy to current immune checkpoint blockades, due to the poor immune activation profiled in NS tumours.

### Activation of the co-stimulatory molecule ICOS increases T cell proliferation in never smoker lung and in the Kras/p53 mouse model of lung cancer

Given the relative quiescence of T cells in early-stage NS tumours, we reasoned that activating co-stimulatory receptors on CD4^+^ T cells may promote more effective anti-tumour responses in these patients than immune checkpoint blockade. We selected ICOS due to its upregulation on tumour infiltrating CD4^+^ T cells in NS tumours compared to other co-stimulatory molecules, albeit at lower levels than that detected in ES patients (Fig 6a,b, Supp Fig 6a). To assess the anti-tumour effects of an anti-ICOS agonist antibody alone or in combination with anti-PD1 blockade *in vivo*, we turned to a genetically engineered model of lung cancer that has low levels of CD8^+^ T cell activation and does not respond to anti-PD1 treatment, the *Kras^G12D/+^;p53^fl/fl^* model^53,54^ (Supp Fig 6b). Similar to our observations in human patients, we found that ICOS expression increased in CD4^+^ T cells in the lungs of *Kras^G12D/+^;p53^fl/fl^* tumour-bearing mice compared with CD4^+^ T cells in control WT mice, making this a suitable model to explore ICOS stimulation *in vivo* (Supp Fig 6c). Tumour burden was unchanged with anti-PD1 or anti-PD1/anti-ICOS combination treatment, yet 2 out 5 mice responded to single agent anti-ICOS treatment (Fig 6c,d). Enhanced CD4^+^ T cell proliferation was observed in the responders in the total lung tissue analysed by flow cytometry (Fig 6e). Spatial analysis by dual immunohistochemistry revealed a greater percentage and number of proliferating CD8^+^ T cells near the small cancerous lesions in the anti-ICOS responders (Fig 6f,g, Supp Fig 6d,e). Interestingly, dual anti-PD1/anti-ICOS treatment was not beneficial for tumour control, perhaps due to enhanced numbers of immune suppressive ICOS^+^PD1^+^ effector T_reg_ cells (Supp Fig 6f,g). T_reg_ cells are potent inhibitors of anti-tumour immunity in the *Kras^G12D/+^;p53^fl/fl^* model^55^, suggesting that anti-ICOS and anti-PD1 dual treatment will need to be evaluated in the context of T_reg_ cell activity. Nevertheless, these data support the notion of employing ICOS agonists to stimulate T cell proliferation and tumour immunity.

**Figure 6.**
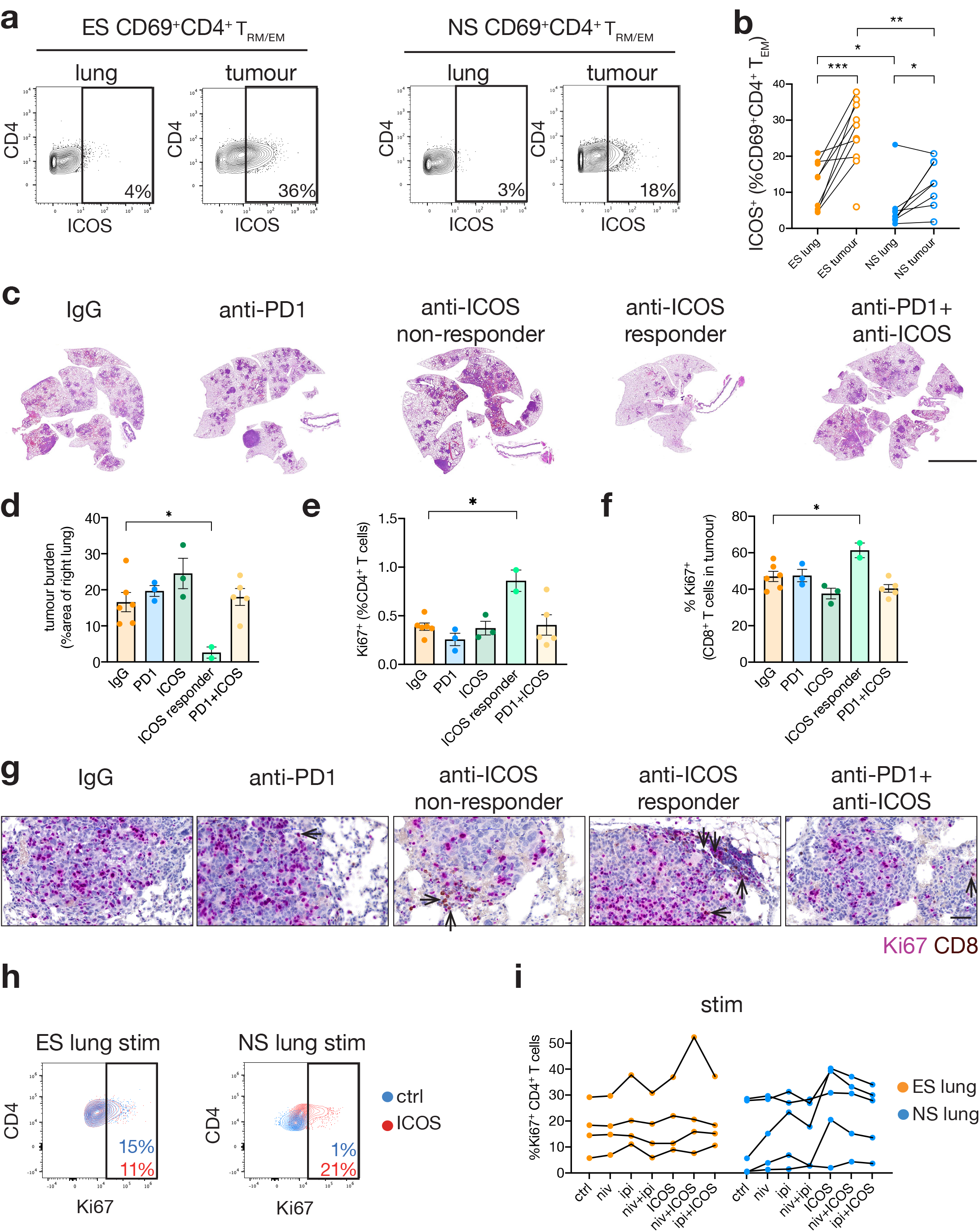
Activation of the co-stimulatory molecule ICOS can enhance T cell proliferation in cancer-associated human and murine T cells. **a**, Representative CyTOF plots of ICOS expression within CD3^+^CD69^+^CD45RO^+^CCR7^-^CD4^+^ T cells in an ES patient (MH0103, 58 yo male, current smoker, stage IIIa LUAD) and NS patient (AH0059, 79 yo female, stage I LUAD). **b**, Quantification of ICOS expression within CD69^+^CD4^+^ T_RM/EM_ in early-stage lung cancers, as analysed by CyTOF. n = 10 ES, n = 8 NS. Paired t-test between lung and tumour, unpaired t-test between non-malignant lung/tumour. *p<0.05, **p<0.01, ***p<0.001. **c**, Representative H&E images of Ad5-CMV-Cre infected *Kras^G12D/+^;p53^fl/fl^* right lung lobes, treated with rat IgG2a and hamster IgG (n = 6), anti-PD1 (n = 3), anti-ICOS (n = 3), anti-ICOS responder (n = 2), or anti-PD1 and anti-ICOS (n = 5). Scale bar indicates 5 mm. **d**, Quantification of tumour burden within right lung lobes in each treatment group. Unpaired t-test, *p<0.05. **e**, Percent of Ki67^+^ cells in CD4^+^ T cells detected by flow cytometry in the left lung lobe. Unpaired t-test, *p<0.05. **f**, Quantification of Ki67^+^CD8^+^ T cells by dual-immunohistochemistry within tumour regions. Unpaired t-test, *p<0.05**. g**, Representative dual Ki67 and CD8 immunostaining within mid-size adenomas in each treatment group. Black arrows indicate dual positive Ki67^+^ and CD8^+^ cells. Scale bar is 50µm. **h,** Representative flow cytometry plots of Ki67 expression in CD4^+^ T cells sorted from non-malignant lung tissue 48h after anti-CD3/anti-CD28 stimulation with anti-ICOS agonist antibody (red) or human IgG4 and hamster IgG-treated control (blue) in IL-2 supplemented media. ES patient (MH0068, current smoker, 68 yo male, LUSC) and NS patient (MH0043, 58 yo female, LUAD). **i**, Quantification of Ki67 expression within CD4^+^ T cells detected by flow cytometry 48h after anti-CD3/anti-CD28 stimulation and treatment with the indicated therapies, compared to human IgG4 and hamster IgG-treated control. n = 4 ES patients, n = 5 NS patients. Unpaired t-test, not significant.

Although tumour tissue was not available, we isolated T cells from non-malignant lung tissue from NS and ES patients to study their potential for responses to an anti-ICOS agonist antibody either alone or together with standard of care checkpoint immunotherapies - nivolumab (anti-PD1 antagonist) and ipilimumab (anti-CTLA4 antagonist). Upon TCR stimulation, treatment with anti-ICOS alone or in combination with nivolumab or ipilimumab enhanced proliferation of CD4^+^ T cells in 3 out of 5 NS patients, and increased Ki67 levels to those detected in cells from ES patients (Fig 6h,i and Supp Fig 6h). These data suggest that ICOS co-stimulation could enhance CD4^+^ T cell activation in NS patients and potentially generate anti-tumour responses.

## Discussion

It is now well accepted that adaptive immunity influences multiple stages of tumour evolution. Cancer immunosurveillance involves T cell elimination of early malignant cells following neoantigen recognition. Later, cancer-immune equilibrium holds premalignant lesions in check. Finally, immune escape involves the development of tumour mechanisms to evade immune predation, enabling invasive lesions to form^2,56^. Yet it remains unclear the extent to which tissue-resident T cells activated by environmental insults may shape the immunogenic evolution of tumours. Our deep profiling of protein markers in non-malignant, early-stage and late-stage primary tumours provides a surrogate for the longitudinal analysis of resident T cells in states of pre-malignancy, early tumour formation and progression. We demonstrate chronic cigarette smoking induces enhanced T_RM_ immunosurveillance in human lungs, yet exerts a potent selective pressure for tumour clones capable of immune evasion early in carcinogenesis. These data identify a critical additional ‘input’ to the system of tumour neoantigen evolution^36^ – the timing of immune pressure – which acts as a major regulator of tumour immune visibility (Figure 7).

**Figure 7.**
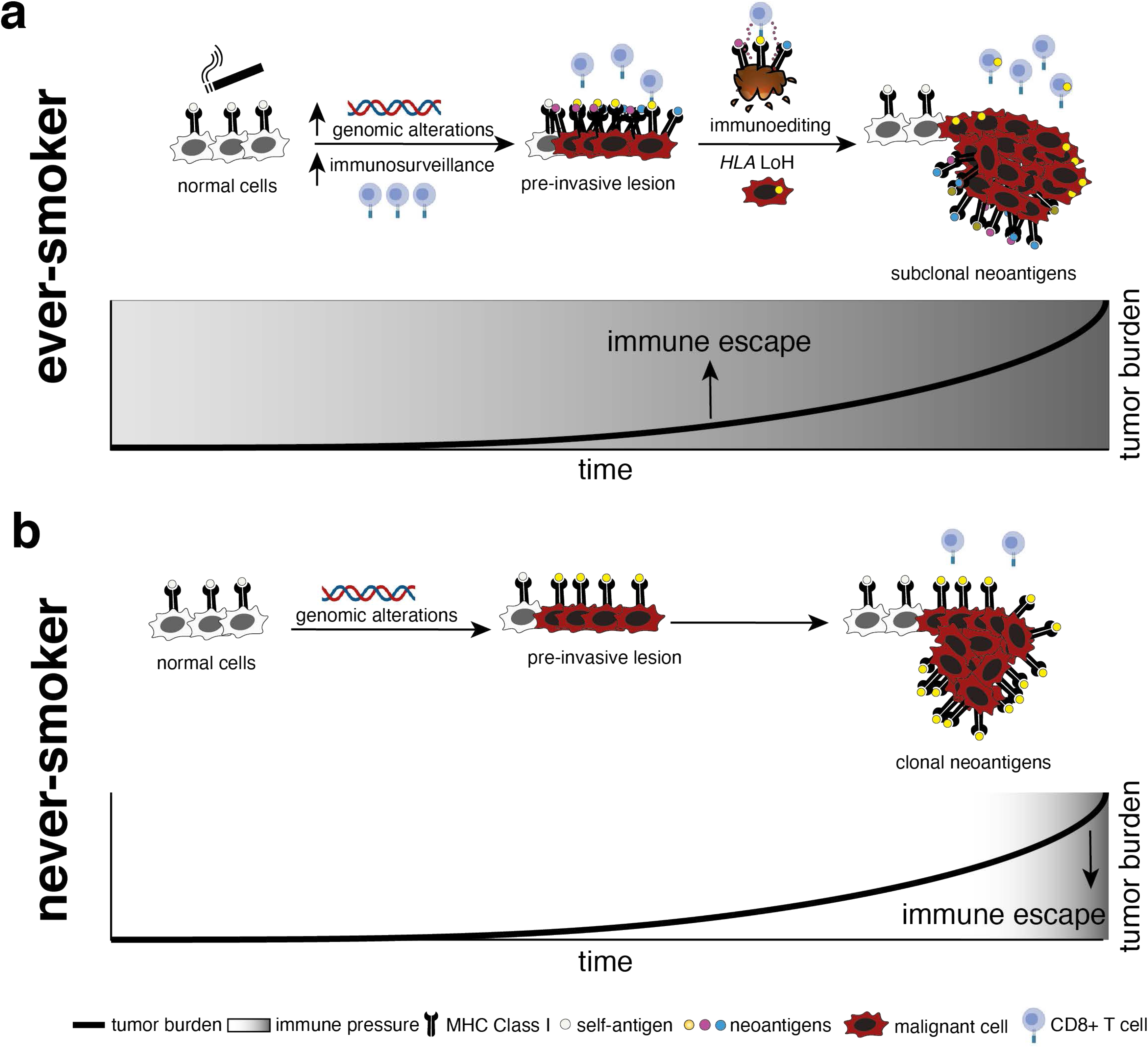
Schematic depicting the timing and intensity of immune pressure and its impact on tumour evolution in ES and NS tumours. **a**, Early immune pressure imposed by smoking activated T_RM_ in the lung of ES patients provoke immune escape mechanisms early in the life of the tumour through *HLA* LoH and neoantigen depletion, resulting in enrichment of subclonal neoantigens in resultant tumours. **b**, Limited immune selective pressure in the early lifespan of NS tumours results in reduced immune escape and visibility to the host T cell response through the expression of clonal neoantigens and intact *HLA*.

T_RM_ in the skin have been shown to be protective against melanoma formation and promote cancer-immune equilibrium^3^. Patients with vitiligo, an autoimmune skin condition characterised by high T_RM_ infiltration, have a lower prevalence of malignant melanoma^57,58^. Further studies in mice demonstrated immunosurveillance by T_RM_ is critical in vitiligo-associated cancer protection, indicating that resident T cell memory is a likely key element in the prevention of tumour onset in the skin^59^. Our discovery of T_RM_ immunosurveillance in ES lungs provides insight into tumour formation in both NS and ES patients. Oncogenic alterations and genomic instability are found in normal lung basal cells from heavy smokers^4,5^, yet not all ever-smokers will develop lung cancers^60^. Pre-invasive LUSC^61^ possess mutations in oncogenic drivers including *KRAS, EGFR* and *KEAP1* but a number of these lesions regress to a normal-like state^62^. The presence of CD8^+^ T cells infiltrating pre-malignant LUSC was proposed to support the elimination of malignant cells^62^. It is tempting to speculate greater T_RM_-immunosurveillance induced by tobacco smoking may protect from invasive cancers induced by this same insult. Evaluating the degree of T_RM_ immunosurveillance within ES and NS pre-invasive LUAD would help resolve this question. Nevertheless, our work suggests that the timing of immune recognition of tumours as influenced by pre-existing immunosurveillance is a critical for resulting tumour immunogenicity.

Early immunosurveillance applies a selective pressure for immune escape mechanisms and has implications for treatment of patients with immune checkpoint inhibitors (ICI). One such escape mechanism involves silencing of the antigen presentation machinery^63^, including loss of MHC Class I by genetic, epigenetic or post-translational modifications. Consistent with previous work^35,64^, we observed that *HLA* LoH occurred irrespective of tumour mutational load. Strikingly, clonal *HLA* LoH was only detected in ES patients in both early-stage and late-stage disease suggesting it is an early event in the evolution of ES tumours. Analysis of *HLA* LoH in preinvasive lung cancer lesions corroborates this observation where *HLA* LoH occurred in 34% of squamous carcinoma *in situ* in ES patients^62^, 8% of adenocarcinoma *in situ*/minimally invasive adenocarcinoma in ES patients and only 1% of these lesions in NS patients^51^. We propose that the T_RM_-rich microenvironment of ES lungs where NSCLC grows is a likely driver of this early immune evasion mechanism. Importantly, *HLA* LoH is a strong predictor of poor response to ICIs in LUAD^64^. Another evasion mechanism is ‘immunoediting’ of tumour cell antigens. As previously reported^37^, immunoediting results in less clonal neoantigens than expected based on the TMB, and we detected this process only in ES patients. A lack of clonal neoantigens also predicts poorer responses to checkpoint blockade^65^. Although ES with NSCLC are thought to be best responders to ICIs, our work provides context to the diversity of responses observed within this group^66^. ES tumours are under immune pressure from their first inception with varied levels of TMB and immune infiltrates among patients. These pressures force early onset of immune escape mechanisms in a subset of patients which ultimately leads to their insensitivity to ICI treatment.

In contrast, NS patients presented as a relatively homogenous group whose tumours evolved in a vacuum of immune selection pressure, resulting in less immune escape of tumour clones. Low T-cell activation and the relatively small numbers of neoantigens diminish chances for T cell recognition, hence generating a *de novo* anti-tumour response with blockade of co-inhibitory molecules has not proved successful for NS patients^12^. However, we propose tumours grown in T_RM_ poor environments, such as those occurring in NS, could paradoxically have favourable features for immune interventions due to the absence of early selective pressure for tumours to become immunologically silent. This lack of immune pressure increases the potential of tumour cell populations to be targeted *en masse* by T cell recognition of clonal neoantigens and intact antigen presentation machinery (Figure 7). Here, the opportunity may lie in the activation of co-stimulatory molecules such as ICOS. Three anti-ICOS humanized antibody agonists are currently in clinical trials in solid tumours, including NSCLC, alone or in combination with anti-PD1 or anti-CTLA4 (NCT02904226; NCT02723955; NCT03251924). Phase I studies have demonstrated no dose-limiting toxicity^67,68^. ICOS agonist treatment will have to be evaluated in the context of the known effect of ICOS on immunosuppressive T_reg_^69^, specifically in combination therapy^70^. Alternative immunotherapeutic approaches for NS patients may also rely on personalized neoantigen vaccines or cell therapy with adoptive transfer of tumour-immunoreactive T cells targeted to these clonal neoantigens^71–74^.

Our work indicates that T_RM_ immunosurveillance induced by cigarette smoking before the onset of malignancy exerts an early selective pressure that acts in addition to tumour cell intrinsic characteristics to shape tumour evolution in lung cancer. Furthermore, this study suggests that T_RM_ present in other solid organs, such as the skin or the breast, might be similarly impacted by environmental insults or physiological dynamics throughout an organism’s lifespan that in turn control cancer immunosurveillance and tumour evolution.

## Methods

### Patient samples

Written informed consent was obtained from all lung cancer patients by the Victorian Cancer Biobank prior to inclusion in the study, according to protocols approved by the WEHI Human Research Ethics Committee (HREC, approval #10/04). Patients were classified as ever-smokers (current and former smokers) or never-smokers (lifetime smoking of less than 100 cigarettes). Matched tumour and adjacent normal lung specimens confirmed by histology were obtained through the Victorian Cancer Biobank from surgically resected tissue of early-stage lung cancer patients (Stage I to IIIa). Primary tumour samples from patients with unresectable, late-stage lung cancers were obtained from endobronchial ultrasound biopsies (EBUS), as was a sample of pulmonary lymph node containing metastatic cancer cells used as a reference control for CyTOF. Patient details are described in Supplementary Table 1. Healthy PBMC used as a reference control was collected from a patient through the Victorian Blood Donor Registry (WEHI HREC approval # 2016.066; Melbourne Health HREC/16/MH/62). Written informed consent was obtained.

### Mice and *in vivo* treatments

All animal experiments were approved by the WEHI Animal Ethics Committee (Approval #2020.001;2020.002;2020.026). Mice were maintained in our animal facilities according to institutional requirements. *Kras^G12D/+^;p53^flox/flox^* mice were obtained from Jackson Laboratory^75^ and maintained on a C57BL/6 background. Equal numbers of male and female mice were intranasally infected with 20 µL of 1 x 10^10^ PFU Ad5-CMV-Cre (University of Iowa Gene Transfer Core) and randomized into treatment groups. Mice were treated with anti-PD1 and anti-ICOS (Supplementary Fig 6b, Supplementary Table 6) 40 days post-infection and collected after 23 days of treatment. Left lung lobes were taken for flow cytometry analyses and right lung lobes were inflated in 4% PFA for histological analyses.

### Human lung and tumour cell preparation

Lung and tumour samples were either processed immediately or held intact for a maximum of 48 h at 4 °C in DMEM/F12 media (Gibco) supplemented with 1 mg/mL of penicillin and streptomycin (Invitrogen). Surgical samples were minced then digested for 1 hour at 37 °C with 2 mg/mL collagenase I (Worthington, #LS004197) and 200 U/mL deoxyribonuclease (Worthington, #LS002140) in 0.2 % D-glucose (Sigma) in DPBS (Gibco). Samples obtained from EBUS biopsies were digested in collagenase/DNase as above for 45 minutes. The cell suspension was filtered through a 100 μm cell strainer and washed with 2% FCS-PBS, followed by red blood cell lysis and further washing with 2%FCS-PBS to obtain a single cell suspension. PBMCs were isolated from whole blood using Ficoll-Paque PLUS (GE Healthcare) separation.

### Mouse lung preparation

Left lung lobes were minced and digested in 2 mg/mL collagenase in 0.2 % D-glucose in DPBS for 45 min at 37 °C. Red blood cells were lysed (0.64% NH_4_Cl) and then cells filtered to obtain a single cell suspension.

### Mass cytometry

Single cell suspensions were pulsed for 1 minute in 25 µM cisplatin (Sigma Aldrich) at room temperature to label dead cells, washed and then fixed in 1.5% PFA (Electron Microscopy Sciences) for 40 min at RT. Cells were then cryopreserved in cell staining media (CSM, PBS with 0.5% BSA and 0.02% sodium azide) and stored at −80°C. Thawed cells were barcoded using a 20-plex palladium isotope barcoding kit (Fluidigm) according to the manufacturer’s directions, blocked with anti-CD16/CD32 FC*γ* II/III (WEHI antibody facility) and stained with extracellular antibodies for 30 minutes at RT (Supplementary Table 2). Cells were permeabilized at 4°C with methanol for 15 minutes, washed thrice in CSM, incubated in 100U/mL heparin for 20 min at RT^76^ and subsequently stained with antibodies against intracellular markers (Supplementary Table 2). Cells were incubated at 4 °C with 125 nM ^191^Ir;^193^Ir DNA intercalator (Fluidigm) in 1.6% PFA overnight and washed in double-distilled water before analysis on a Helios CyTOF (Fluidigm, maintained by Materials Characterisation and Fabrication Platform, University of Melbourne). Antibody conjugates that could not be purchased were created from carrier-free antibody solutions conjugated to custom isotopes (Trace Sciences, Fluidigm) using the Maxpar kit (Fluidigm). Antibodies were validated (Supp Fig 1a-c) and titrated to ensure specificity and sensitivity. Patient samples were analysed in batches, with equal numbers of ES and NS patients per run and the inclusion of two reference controls (PBMC and an EBUS metastatic lymph node biopsy) per run to detect batch-to-batch variability.

### Analysis of mass cytometry data

All .fcs files generated were concatenated, normalized to beads and debarcoded using the R package Catalyst^77^ or Premessa. The dataset was further batch-corrected using the CytofRUV R package^39^ and a k value of 2 using two reference control samples (PBMC and metastatic lymph node) that were contained in each batch analysed. Normalised data was then subject to FlowSOM clustering^78^ and filtering to identify CD45+ cells using cell classification markers (Supp Table 2, 6 CD3^+^ clusters identified and 8 CD3^-^ clusters identified) and visualised using t-SNE. CD4+ and CD8+ T cells were combined across patients and subjected to viSNE^79^ analyses using CytoBank software. viSNE analyses were conducted on all phenotyping markers (Supp Table 2) and subject to equal scaling. Samples were also subject to manual gating using reference control samples to standardize between runs (FlowJo) and the R packages limma/pheatmap^41^ used to analyse the data expressed as a proportion of the parent gate.

### Flow cytometry

To isolate human T cells from non-malignant lung tissues, single cell suspensions were blocked with anti-CD16/CD32 FC*γ* II/III (WEHI antibody facility) before staining with anti-CD235a-PE, anti-CD140b-PE, anti-CD31-PE, anti-CD45-BV510, anti-CD4-PerCP-Cy5.5, anti-CD8-Pacific Blue (Supplementary Table 3). Cells were washed and resuspended in 0.5 µg/mL propidium iodide. CD45+CD4+CD8+ cells were sorted using a 100 µm nozzle on either a FACSAria or FACSAria Fusion (BD) and processed immediately after.

For intracellular staining of mouse and human cells, single cell suspensions were incubated with fixable live/dead GREEN (Invitrogen) or Zombie Aqua (Invitrogen) according to the manufacturer’s instructions to distinguish viable cells. Cells were washed with 2%FCS-PBS, blocked with anti-CD16/CD32 FC*γ* II/III (WEHI antibody facility) and stained with extracellular antibodies for 30 min at 4 °C. Cells were fixed and permeabilised with Foxp3/Transcription Factor Staining Kit (eBiosciences) and stained with intracellular antibodies for 30 min at 4 °C (Supplementary Table 3 and Supplementary Table 5). Samples were acquired on Aurora spectral unmixing cytometers (CyTEK) and analysed using Flowjo and Cytobank.

### Human lung T cell *in vitro* assays

96-well round bottom plates were coated with 5µg/mL anti-CD3 for a minimum of 2 hours at 37 °C before rinsing and removing. Human T cells were plated in stimulated (anti-CD3 and with 1µg/mL anti-CD28) or unstimulated conditions in IMDM media supplemented with 10% fetal calf serum, 1% HEPES, non-essential amino acids, glutamax, sodium pyruvate (all Gibco), 50 µM 2-mercaptoethanol (Sigma) and 100U/mL recombinant IL-2 (Peprotech). Immunotherapeutics were added detailed in Supplementary Table 6. Cells were collected for flow cytometry analysis 48 and 72 hours after drug addition, Golgi Stop and Golgi Plug (BD) were added to the culture media 3 hours before each collection.

### Immunohistochemistry

Human tissue was formalin fixed and paraffin embedded, before antigen retrieval with citrate buffer (10mM, pH 6) and blocking with 10% goat serum. Sections were incubated with antibodies against CD45 or MHC Class I for 30 mins at RT (Supplementary Table 7) and stained with biotinylated anti-mouse secondary antibody (Vector Lab) before counterstaining with haematoxylin. Mouse tissue was paraformaldehyde fixed and paraffin embedded, before low pH antigen retrieval and staining with CD8a and Ki67 (Supplementary Table 7) antibodies and secondary antibody (EnVision+ HRP-rabbit, Dako, catalog #K400311-2) using the EnVision DuoFlex system (Dako). CD8a was detected with DAB (Dako) and Ki67 was detected with Magenta Substrate Chromogen (Dako).

### Spatial analysis of dual CD8/Ki67 staining

Cell detection, cell counting, and analysis of marker positivity was performed in an automated manner using QuPath^80^. Colour deconvolution was first completed to separate hematoxylin from different markers. Cells were detected with Stardist^81^, an object detection tool using machine learning. To measure whether a cell was positive to a certain marker, an automated method was used. The cutoff value was based on the distribution of mean intensities per marker. Considering positive cells as outliers, we assigned a cell to be positive to a marker if the mean intensity relative to that marker was more than 4 median absolute deviations above the median. Subsequently, we assigned each cell to be Ki67^+^ only, CD8^+^ only, CD8^+^ Ki67^+^ double positives or unidentified. For the area measurement, we first used a multilayer perception neural network (MLP) as a pixel classifier in order to differentiate tumour from non-tumour areas. Ten regions per class (tumour, non-tumour) were manually annotated for the training. The tumour regions were then identified on the whole tissue using the pixel classifier. Finally, CD8^+^ Ki67^+^ double positive cell densities were computed in tumour areas.

### Patient datasets

Access to the TRACERx cohort of 100 NSCLC patients with 327 tumor regions and matched germlines^82^ was granted by a Data Access Agreement between the Francis Crick Institute and the Walter and Eliza Hall Institute of Medical Research (DAC #2020-0132). Whole-exome sequencing (WES) alignment files were obtained from the European Genome-phenome Archive (dataset EGAD00001003206). Clinical data and somatic variants were extracted from Tables S2 and S3 in Supplementary Appendix 2 and smoking signatures from Supplementary Appendix 1 of Jamal-Hanjani *et al*^82^. Variants were converted to standard VCF files for downstream analyses.

Access to the LxG cohort of 20 lung cancer patients with 39 tumor regions and matched germlines^83^ was granted by the Garvan Institute of Medical Research. Whole-genome sequencing (WGS) alignment files, somatic variants, cellularities, smoking signatures and clinical data were obtained from the authors and downloaded from the National Computational Infrastructure of Australia. We excluded 1 patient and 4 tumor regions from analysis due to low cellularity (<20%).

Access to the Chen cohort of pre-invasive (n = 98, adenocarcinoma in situ, minimally invasive adenocarcinoma) and invasive LUAD (n = 99) was granted by the FUSCC Chen group. Whole transcriptome (RNAseq) alignment files were obtained from the European Genome-phenome Archive (dataset EGAD00001005479). Clinical data, TMB and HLA LoH were extracted from the publicly available Source Data of Chen *et al*^51^.

We used publicly available data from the TCGA cohort of LUAD^7^, using TMB calculated from Hoadley et al^84^ and neoantigens from Rooney et al^50^. All datasets are detailed in Supplementary Table 8.

### Tumour mutational burden

Tumor mutational burden (TMB) was calculated for all samples in the LxG and TRACERx by dividing the number of somatic variants passing variant caller quality thresholds by the total capturable area (3,200Mb for LxG and 50Mb for TRACERx).

### Immune infiltrate estimates

Immune infiltrate estimates from RNAseq of TRACERx were used from Rosenthal *et al.*^37^. Immune infiltration for the TCGA LUAD cohort was estimated using the TIMER2.0 tool^85^. Immune infiltration for the Chen cohort was estimated by first generating read counts and transcript lengths per gene using featureCounts (v1.6.3) ^86^, then TPM-normalising them using ADImpute (v 0.99.3) and finally using the TIMER2.0 tool^85^.

### HLA typing and LoH calling

Haplotyping of *HLA-A*, *HLA-B* and *HLA-C* was carried out on all germline and tumor samples in both cohorts using POLYSOLVER (v4) with default parameters^87^. Loss of heterozygosity (LoH) was inferred for each patient by comparing HLA types in tumor samples against the germline type at the resolution of allotype and subtype (2 sets of digits). POLYSOLVER allowed for the analysis of LoH in both WES and WGS datasets. The loss of at least one allele classified the sample as affected by HLA LoH. Clonal LoH was called when an allele was lost in all samples for a patient.

### Neoantigen predictions

Neoantigen prediction was carried out using pVACseq (v3.0.3-1) within the pVACtools suite via Docker^88^. Variants were annotated using Variant Effect Predictor^89^ with the Wildtype and Frameshift plugins, to produce mutant and wild type peptides per variant, as required by pVACseq. HLA types and variants per sample were entered into pVACseq to predict neoantigens and simultaneously account for loss or gain of HLA alleles in tumors. Neoantigens were predicted for peptides 8-10 amino acids in length where a binding affinity of IC_50_<500 nM was obtained in the mutant peptide sequence against any of the sample’s HLA types. The coverage filter was modified to require a tumor variant allele frequency of >0.06 and tumor depth of >40 to allow the analysis of low cellularity samples with high sequencing quality. All other parameters used default values. Summary counts and clonality of neoantigens were determined using a custom script that parsed the filtered, condensed and ranked output files from pVACseq. Clonal neoantigens were called as neoantigens present in all samples from a particular patient.

### Immunoediting

The power transformed square root of both TMB and neoantigen counts were calculated per sample in the TRACERx and LxG cohorts to account for an inability to log transform zeros:

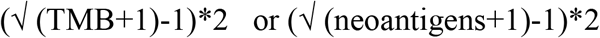

Samples were then plotted in R and using the rlm function was used to fit a robust linear model, the weight and residuals of each datapoint were calculated. Samples with weights < 0.9 and negative residuals were considered ‘immunoedited’ – ie, those having fewer than expected numbers of neoantigens for a given TMB. Other approaches in the literature to calculate expected neoantigen numbers for a given TMB from a reference dataset e.g. TCGA, are subject to data quality issues, the influence of mutational spectra and a lack of patient germline HLA types^34^.

## Acknowledgements

We thank Dr Alexandra Garnham for advice on analysing CyTOF data, Alissa Robbins and Tania Tan for T cell reagents and CyTOF advice, Dr Sarah Best for mouse model advice and critical reading of the manuscript. We thank WEHI facilities including Histology (Emma Pan and Ellen Tsui), Flow cytometry (Simon Monard) and Bioservices (Shannon Oliver) for expert advice and for performing experiments. We are grateful to the Victorian Cancer Biobank and all lung cancer patients who participated in this study. This work was performed in part at the Materials Characterisation and Fabrication Platform (MCFP) at the University of Melbourne and the Victorian Node of the Australian National Fabrication Facility (ANFF), with support from the Victorian Comprehensive Cancer Centre under the Resistance to Targeted Therapies Program. The mass cytometry studies were also supported in part by the Australian Cancer Research Foundation. C.W. is supported by a Deep Machanda Lung Foundation Australia Post-Doctoral Fellowship and a Cure Cancer grant. C.L.G. is supported by funding from a NHMRC Early Career Fellowship (GNT1160963). D.H.D.G is supported by Australian NHMRC Fellowships/grants (1090236, 1158024 and 1145888), Cancer Council of Victoria Grants-in-Aid (1146518 and 1102104). M.L.A.L. is supported by funding from the Viertel Foundation Senior Medical Research Fellowship. This work is supported by a NHMRC Ideas Grant (GNT1182155), The Harry Secomb Trust, the Jenny Tatchell Fund and by funds from the Operational Infrastructure Support Program provided by the Victorian Government and NHMRC IRIISS (Independent Research Institutes Infrastructure Support Scheme) Grant.

---

**Supplementary Figure 1. Validation of antigen detection by CyTOF conjugates. a,** Representative antibody validation experiments for the specificity of epithelial markers (Sox2) in LUSC and LUAD cell lines (left), cell death markers (cleaved caspase 3) in a LUSC cell line treated with BH3 mimetics (A1331852: BCL-XL inhibitor; S63845: MCL1 inhibitor) to induce apoptosis^91^ (middle) and T cell co-stimulatory markers (OX40) in human CD4^+^ T cells from bulk PBMCs stimulated for 48h with anti-CD3/anti-CD28. **b**, Quantification of percentage of CD45^+^ cells in non-malignant lung and tumour tissue from ES and NS patients by CyTOF (left panel) and immunohistochemistry (right panel, representative patients). CyTOF CD45^hi^ defined as >80% CD45^+^ cells; CyTOF CD45^lo^ defined as <20% CD45^+^ cells. n = 10 ES, n = 8 NS. Paired t-test between lung and tumour, unpaired t-test between non-malignant lung/tumour. *p<0.05. **c**, Correlation between PDL1 expression in pan-keratin^+^ tumour cells detected by CyTOF (y-axis) and PDL1 expression detection by clinical IHC, expressed as tumour proportional score. n = 9 early-stage primary tumours with matched CyTOF and clinical data available. **d**, Heatmap of the median cell type marker expression (scaled, arcsinh-transformed) across all samples and aggregated by clusters identified in Fig 1a. **e**, Quantification of individual cell cluster in non-malignant lung tissue and tumour tissue determined by FlowSOM clustering, expressed as a percentage of CD45^+^ cells. Paired Wilcoxon ranked tests between lung and tumour, unpaired t-test between samples from patients with different smoking statuses, p-value not corrected for multiple testing. *p<0.05, **p<0.01.

**Supplementary Figure 2. Phenotype and activity of tissue resident memory T cells within non-malignant lung tissue of ES and NS lung cancer patients. a**, Heatmap of the proportion of co-stimulatory and co-inhibitory receptor expressing cells within CD3^+^CD69^+^CD45RO^+^CCR7^-^ resident T_reg_ (left) and CD8^+^ T_RM_ cells (right) in non-malignant lung tissue analysed by CyTOF. Data is scaled to mean expression ± standard deviation per marker. N = 9 ES and n = 10 NS patients. Unpaired t-test, not significant. Greyed out bars indicate patients with too few cells (<50) to reliably assess cell phenotypes.

**Supplementary Figure 3. Proportions of T cells infiltrating NS and ES tumours are variable. a**-**d,** Proportion of CD69^+^CD45O^+^CCR7^-^ T_RM_-like cells in CD3^+^ T cells. Total T_RM_-like cells (**a**) CD4^+^ T_RM_-like cells in CD4^+^ T cells (**b**), resident-like activated T_reg_ in CD4^+^ T cells (**c**), and CD8^+^ T_RM_-like cells in CD8^+^ T cells (**d**). Paired t-test between lung and tumour, unpaired t-test between NS and ES. p-values not corrected for multiple testing. *p<0.05.

**Supplementary Figure 4. Relationship between tumour mutational burden, neoantigen numbers, CD8^+^ T cell infiltration and smoking status in lung adenocarcinoma. a**, Association between TMB and neoantigen numbers in the TCGA LUAD cohort, coloured according to smoking status. Linear regression analysis, p value as indicated. **b**, TMB in the TCGA LUAD cohort related to patient smoking status. Unpaired t-test with Welch’s correction, ***p<0.001. **c**, Number of neoantigens in TCGA LUAD cohort related to patient smoking status. Unpaired t-test with Welch’s correction, ***p<0.001. **d**, TMB in the TRACERx cohort of LUAD, NSCLC-NOS and LUSC related to patient smoking status. Unpaired t-test with Welch’s correction, ***p<0.001. Data represents multiple samples per indicated numbers of patients. **e**, Number of neoantigens in the TRACERx cohort of LUAD, NSCLC-NOS and LUSC related to patient smoking status. Unpaired t-test with Welch’s correction, ***p<0.001. Data represents multiple samples per patient. **f**, Scatter plot of TMB and CD8^+^ T cell score in the TCGA LUAD cohort. TMB calculated from whole exome sequencing data^84^ and CIBERSORT CD8^+^ T cell score estimated from RNA sequencing data using TIMER2.0 online platform^85^. Data represents the power transformed square root of values to account for zero values not possible using log transformation. Linear regression analysis, p value as indicated. **g**, Scatter plot of neoantigen numbers and CD8^+^ T cell score within the TCGA LUAD cohort. Neoantigens calculated from whole exome sequencing data as published in Rooney *et al*^50^. Linear regression analysis, p value as indicated. **h,** Volcano plot depicting fold change in T_RM_ accessory molecules between NS and ES late-stage tumours. 30 parameters analysed by CyTOF, co-stimulatory molecules in green, CTLA4 in yellow and co-inhibitory markers in purple. Dotted line indicates q value = 0.05. Data represents average marker fold change within each cell type (CD4^+^ T_RM_-like, resident-like Treg, and CD8^+^ T_RM_-like), n = 6 ES and n = 4 NS late-stage (Stage IIIb/IV) tumours.

**Supplementary Figure 5. Clonal immune escape events are only detected in ever-smoker patients. a**-**c**, Proportion of patients with loss of heterozygosity of one or more *HLA* alleles in some tumour sites (subclonal LoH; green) or all sites (clonal LoH; blue) in the TRACERx cohort of LUAD, NSCLC-NOS and LUSC related to smoking status (**a**), TMB (**b**) or immune infiltration estimated from RNA sequencing data (**c**). Fisher’s exact test, *p<0.05. **d**, *HLA* LoH in adenocarcinoma *in situ* and minimally invasive adenocarcinoma, the pre-invasive forms of LUAD related to smoking status, data called from Chen *et al*^51^. Clonality of LoH cannot be determined from single site data. Fisher’s exact test, not significant. **e**, TMB and the number of neoantigens calculated in samples from the TRACERx and LxG cohort were subject to robust linear modelling and residuals and weights of each data point are plotted. Samples with a weight < 0.9 and a negative residual are indicated in red as ‘immunoedited’ or neoantigen depleted. **f**, Proportion of patients with clonal immunoediting (present in all sites) or subclonal immunoediting (present in some sites) in the TRACERx cohort of LUAD, NSCLC-NOS and LUSC, related to TMB, immune infiltration, HLA LoH or smoking status. Fisher’s exact test, *p<0.05. **g**, Proportion of neoantigens shared in all tumour sites (clonal neoantigens) in the TRACERx cohort of LUAD, NSCLC-NOS and LUSC, related to TMB, immune infiltration, *HLA* LoH events and immunoediting. Unpaired t-test, **p<0.01.

**Supplementary Figure 6. Targeting anti-ICOS as an alternative immunotherapy in T cell quiescent lung tumours. a**, Quantification of co-stimulatory molecule OX40 and 4.1BB expression in CD69^+^CD45RO^+^CCR7^-^ CD4^+^ T cells in early-stage lung cancers, as analysed by CyTOF. n = 10 ES, n = 8 NS. Paired t-test between lung and tumour, unpaired t-test between non-malignant lung/tumour. P-values not corrected for multiple testing, *p<0.05, **p<0.01. **b**, Schematic of *in vivo* treatment of Ad5-CMV-Cre intranasally infected mice bearing *Kras^G12D/+^;p53^fl/fl^* tumours with either rat IgG2a and hamster IgG control; single agent anti-PD1 and hamster IgG, single agent anti-ICOS agonist antibody and rat IgG2a or combination anti-PD1 and anti-ICOS. **c,** Expression of ICOS in CD4^+^ T cells in the spleen and left lung lobe of *Kras^G12D/+^;p53^fl/fl^* mice, tumour-free (Ad-CMV-Cre -) or in tumour-bearing lungs (intranasal infections with Ad-CMV-Cre +) measured by flow cytometry. **d**, Representative images of spatial quantification of Ki67^+^(magenta substrate chromogen, MSC) CD8^+^ (DAB) cells in tumour areas using QuPath software. Scale bar indicates 20 µm. **e**, Density of Ki67^+^CD8^+^ T cells per mm^2^ of tumour in each treatment group. **f**, Number of ICOS^+^CD62L^-^ effector T_reg_ cells^92^ in the T_reg_ compartment in the left lung lobe as detected by flow cytometry in each treatment group. Unpaired t-test, *p<0.05. **g**, Proportion of PD1^+^ cells in the effector T_reg_ population in the left lung lobe as detected by flow cytometry in each treatment group. Unpaired t-test, *p<0.05, **p<0.01. **h,** Percent of Ki67^+^ cells in CD8^+^ T cells detected by flow cytometry 48h after anti-CD3/anti-CD28 stimulation and treatment with the indicated therapies. Control (ctrl) is human IgG4 and hamster IgG treatment. n = 4 ES patients, n = 5 NS patients. Unpaired t-test, not significant.

