## Supplemental Figures and Tables for "Early immune pressure imposed by tissue resident memory T cells sculpts tumour evolution in non-small cell lung cancer"

Weeden *et al*, Supp Figure 1

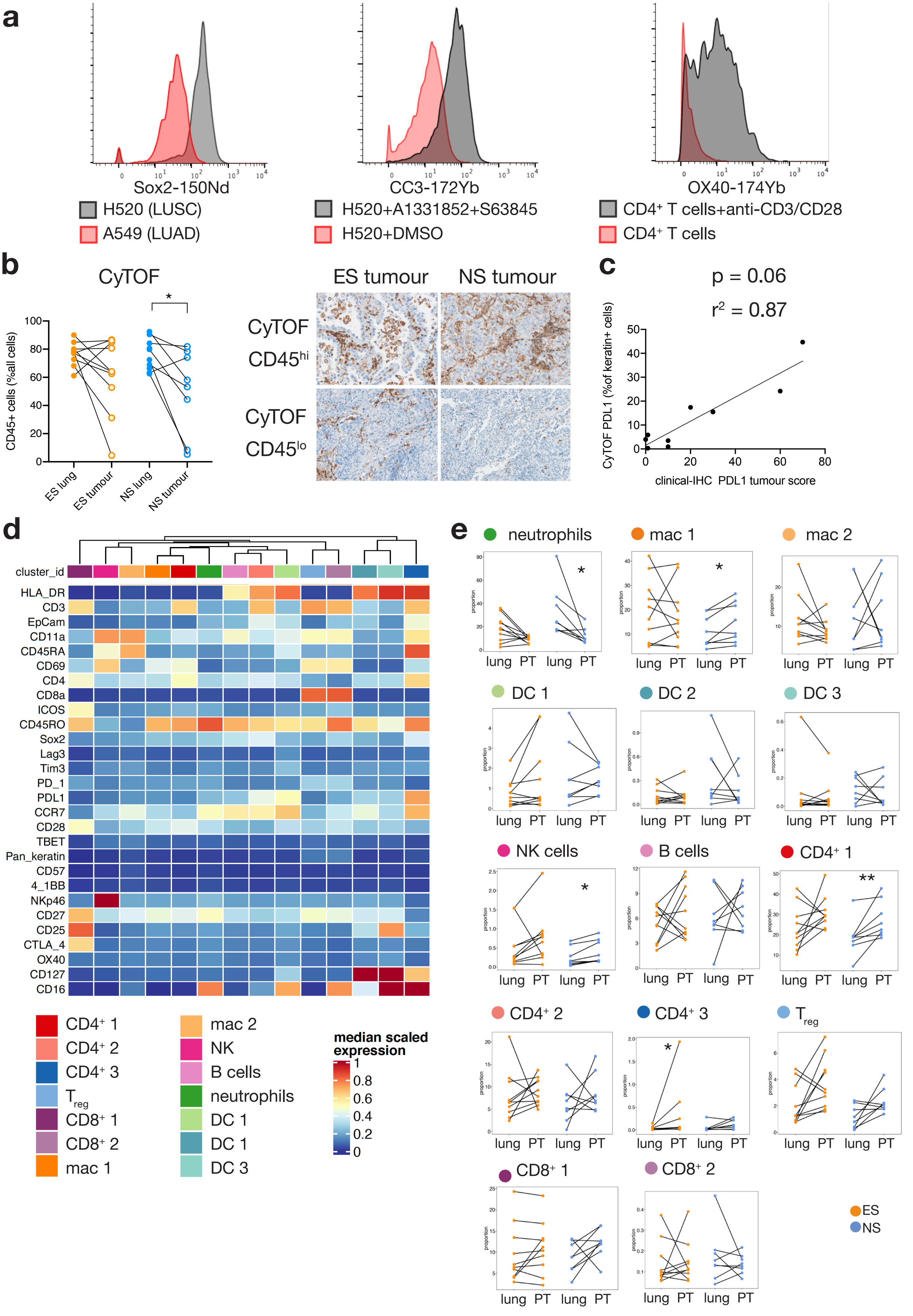

**a**

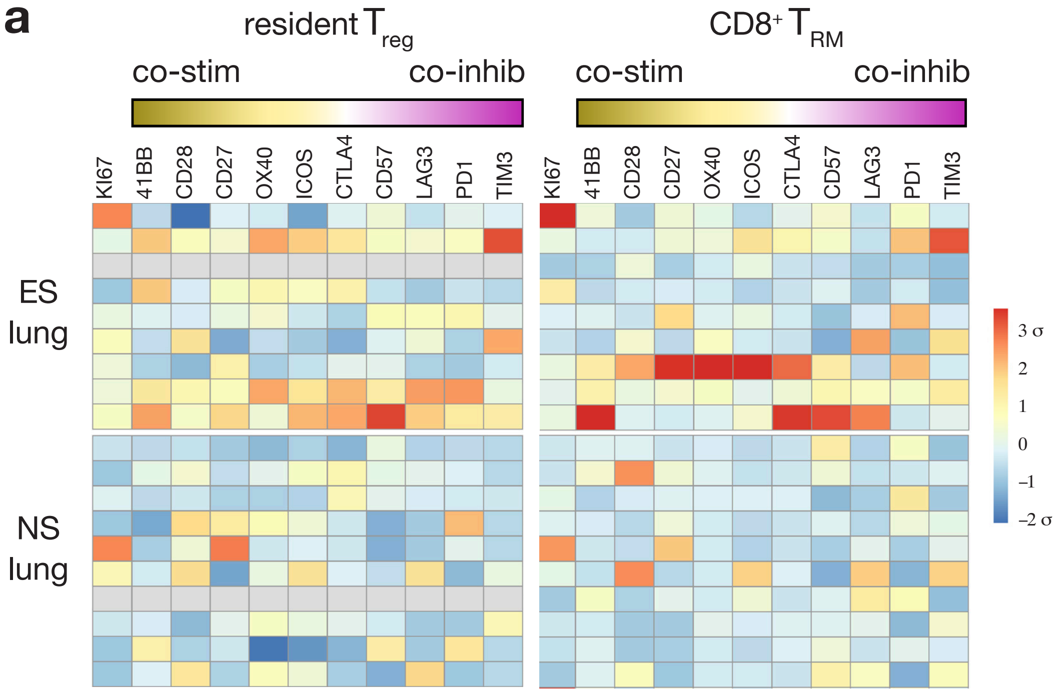
resident $T_{reg}$
co-stim
co-inhib
$CD8^{+}$ $T_{RM}$
co-stim
co-inhib
KI67
41BB
CD28
CD27
OX40
ICOS
CTLA4
CD57
LAG3
PD1
TIM3
ES lung
NS lung
3 σ
2
1
0
–1
–2 σ

Weeden *et al*, Supp Fig 3

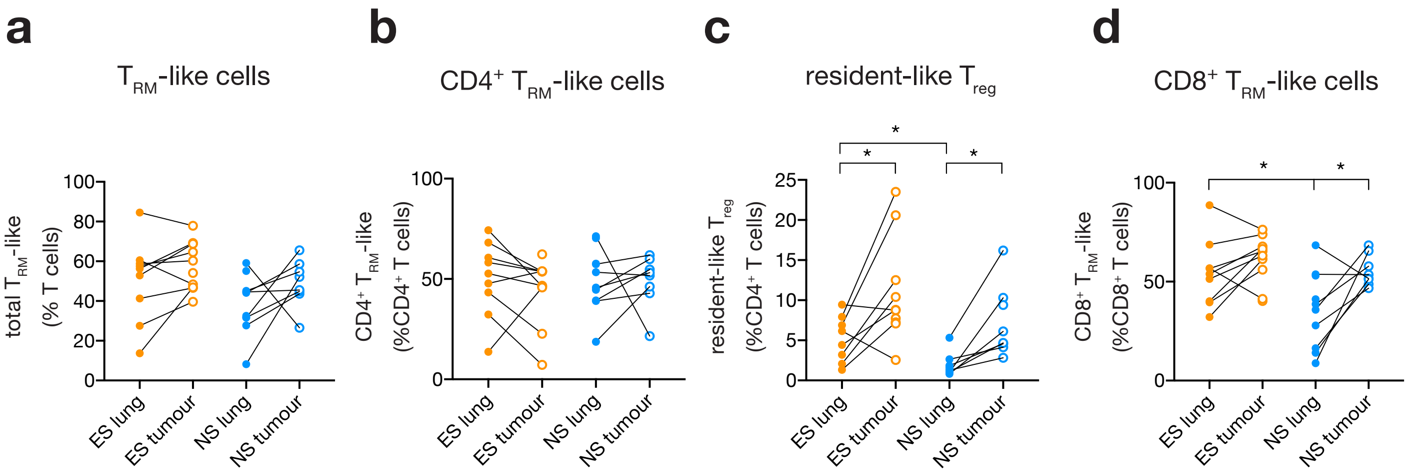

Weeden *et al*, Supp Figure 4

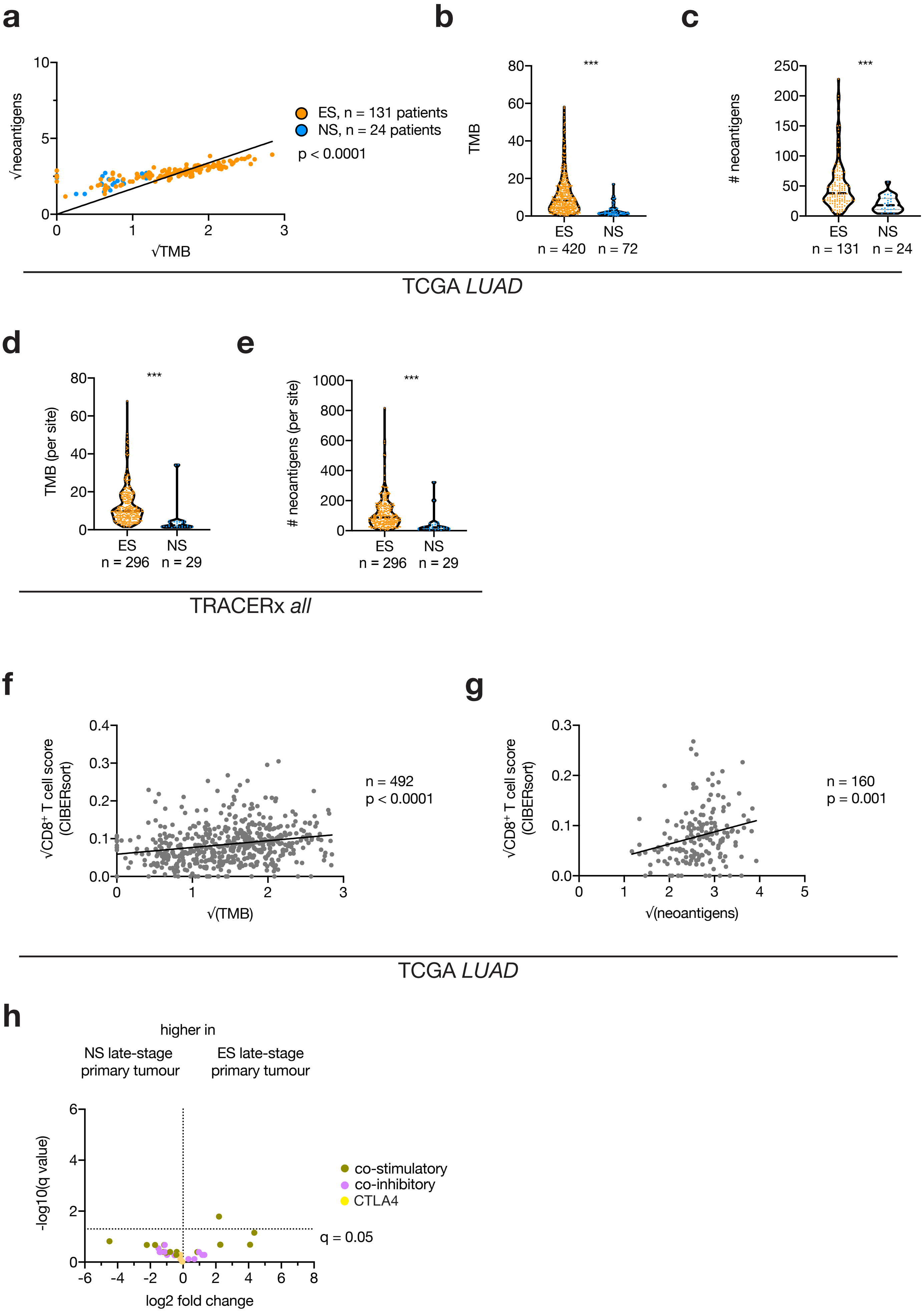

Weeden *et al*, Supp Figure 5

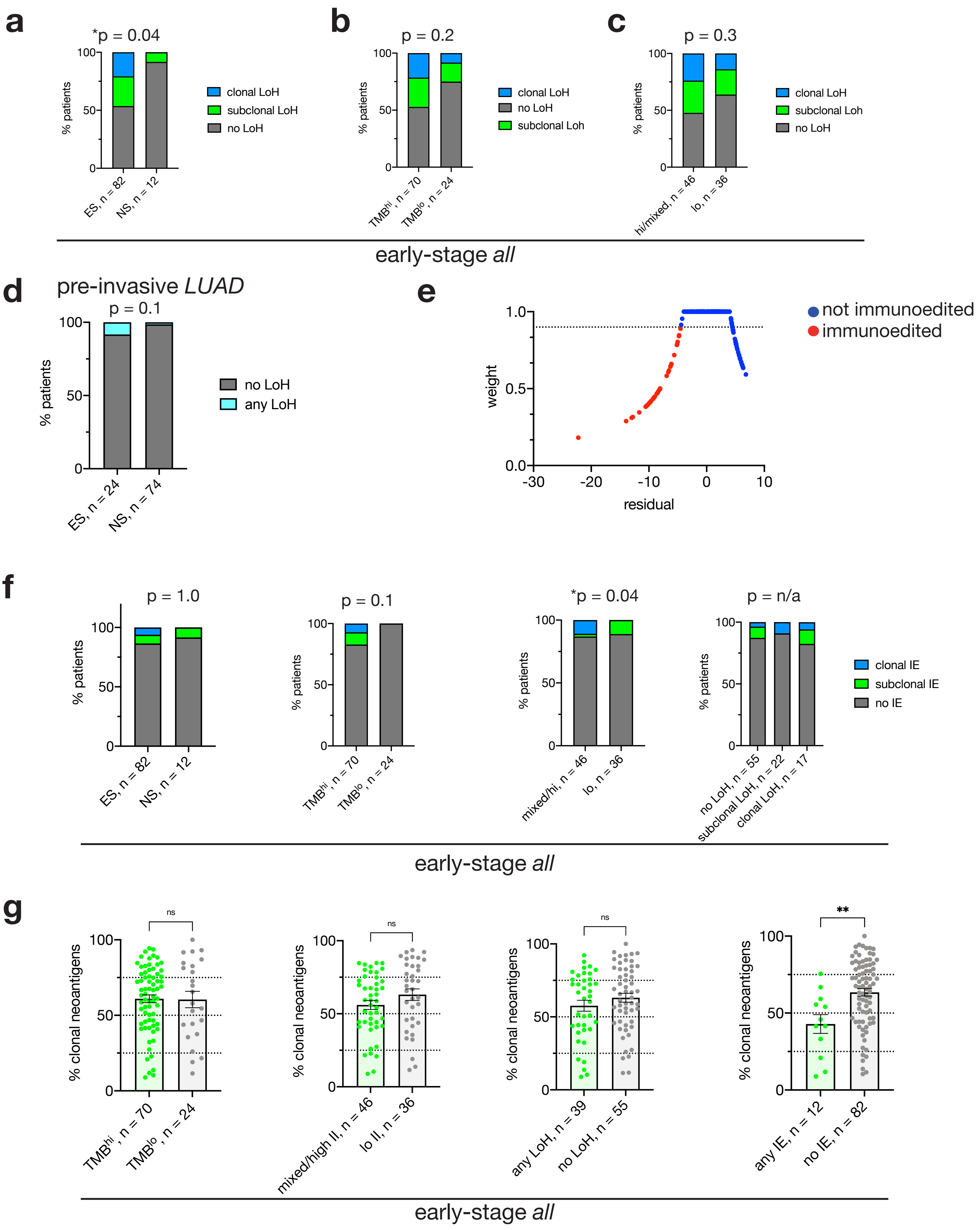

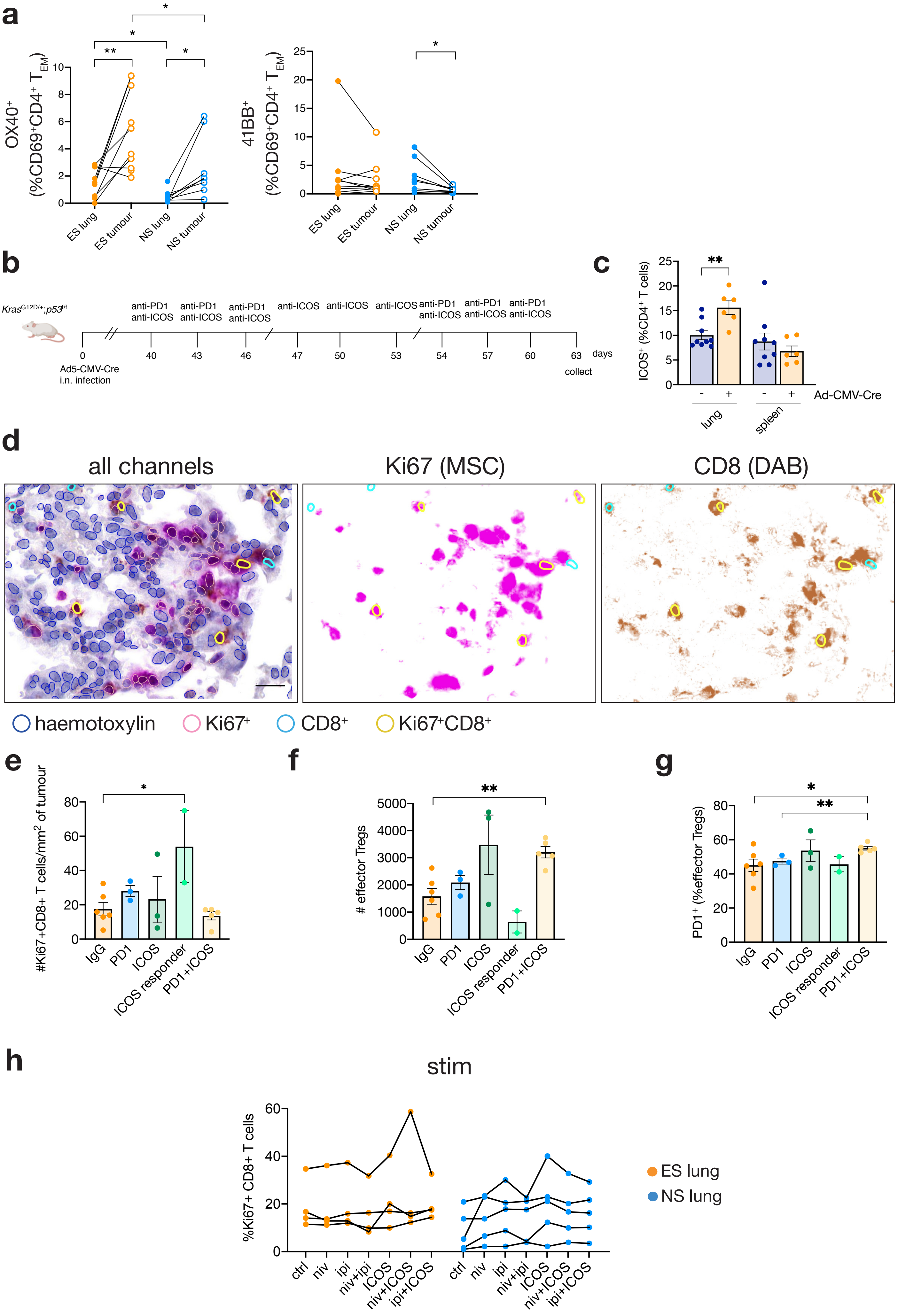

a
OX40+ (%CD69+CD4+ TEM)
ES lung
ES tumour
NS lung
NS tumour
41BB+ (%CD69+CD4+ TEM)
b
KrasG12D/+;p53f/f
anti-PD1 anti-ICOS
anti-ICOS
days
Ad5-CMV-Cre i.n. infection
collect
c
ICOS+ (%CD4+ T cells)
Ad-CMV-Cre
lung
spleen
d
all channels
Ki67 (MSC)
CD8 (DAB)
haemotoxylin
Ki67+
CD8+
Ki67+CD8+
e
#Ki67+CD8+ T cells/mm2 of tumour
f
### effector Tregs
g
PD1+ (%effector Tregs)
IgG
PD1
ICOS
ICOS responder
PD1+ICOS
h
stim
%Ki67+ CD8+ T cells
ctrl
niv
ipi
niv+ipi
ICOS
niv+ICOS
ipi+ICOS
ES lung
NS lung

Weeden et al, “Early immune pressure imposed by tissue resident memory T cells sculpts tumour evolution in non-small cell lung cancer”

| *early-stage lung and lung tumours analysed by mass cytometry* | | |
|---|---|---|
| | **ever-smokers n=10** | **never-smokers n=10** |
| sex | 6M, 4F | 2M, 8F |
| age (mean, range) | 66.7 (47 – 83) | 64.9 (47 – 79) |
| smoking history | 6 ex-, 4 current-smokers | 10 never-smokers |
| *lung cancer subtype* | | |
| adenocarcinoma | 7 | 8 |
| squamous cell carcinoma | 3 | - |
| carcinoid | - | 1 |
| other | - | 1 |
| *stage* | | |
| I | 2 | 3 |
| II | 5 | 4 |
| IIIa | 3 | 2 |
| IV | - | - |
| n/a | - | 1 |
| *late-stage patients analysed by mass cytometry* | | |
| | **ever-smokers n=7** | **never-smokers n=5** |
| sex | 3M, 4F | 3M, 2F |
| age (mean, range) | 66 (59 – 78) | 67 (55 – 76) |
| smoking history | 2 ex-, 5 current-smokers | 5 never-smokers |
| *lung cancer subtype* | | |
| adenocarcinoma | 2 | 4 |
| squamous cell carcinoma | 1 | - |
| carcinoid | - | - |
| NSCLC-NOS | 4 | 1 |
| other | - | - |
| *stage* | | |
| I | - | - |
| II | - | - |
| III | 5 | 2 |
| IV | 2 | 3 |
| n/a | - | - |
| *early-stage lung analysed by flow cytometry and/or in vitro T cell assays* | | |
| | **ever-smokers n=6** | **never-smokers n=5** |
| sex | 2M, 4F | 2M, 3F |
| age (mean, range) | 62.2 (47 – 80) | 57.2 (47 – 77) |
| smoking history | 2 ex-, 4 current-smokers | 5 never-smokers |
| *lung cancer subtype* | | |

| | | |
|---|---|---|
| adenocarcinoma | 5 | 4 |
| squamous cell carcinoma | 1 | - |
| carcinoid | - | 1 |
| other | - | - |
| *stage* | | |
| I | 2 | 4 |
| II | 4 | 1 |
| IIIa | - | - |
| IV | - | - |
| n/a | - | - |

**Supplementary Table 1:** First panel contains patient details for non-malignant lung and early-stage tumour analyses by mass cytometry. All tissue acquired from surgical resections. Second panel contains patient details for late-stage primary tumour analysed by mass cytometry, all tissue acquired from endobronchial ultra-sound guided biopsies. Third panel contains patient details for non-malignant lung tissue analysed by flow cytometry and subject to *in vitro* T cell assays. All tissue acquired from surgical resections for lung cancer, tumour specimens unavailable under COVID-19 protocols.

| *human CyTOF antibodies* | | | | | |
|---|---|---|---|---|---|
| *cell classification* | | | | | |
| **antigen** | **clone** | **conjugate** | **manufacturer** | **catalog #** | **dilution** |
| EpCam | 9C4 | 141 Pr | Fluidigm | 3141006B | 1/100 |
| Pan keratin | C11 | 162 Dy | Fluidigm | 3162027A | 1/100 |
| p63* | EPR5701 | 165 Ho | Abcam | ab214790 | 1/25 |
| TTF1* | EP1584Y | 173 Yb | Abcam | ab186822 | 1/100 |
| HLA-A,B,C* | W6/32 | 150 Nd | WEHI | n/a | 1/100 |
| CD45 | HI30 | 89 Y | Fluidigm | 3089003B | 1/100 |
| CD11a | H111 | 142 Nd | Fluidigm | 3142006B | 1/100 |
| CD16 | 3G8 | 209 Bi | Fluidigm | 3209002B | 1/100 |
| NKp46* | 9E2 | 166 Er | Biolegend | 331902 | 1/50 |
| HLA-DR* | L243 | In115 | Biolegend | 307602 | 1/50 |
| CD3* | UCHT1 | 139 La | Biolegend | 300443 | 1/50 |
| CD4 | RPA-T4 | 145 Nd | Fluidigm | 3145001B | 1/100 |
| CD8a | RPA-T8 | 146 Nd | Fluidigm | 3146001B | 1/100 |
| CD127 | A019D5 | 176 Yb | Fluidigm | 3176004B | 1/100 |
| CD25 | 2A3 | 169 Tm | Fluidigm | 3169003B | 1/100 |
| CD45RO | UCHL1 | 149 Sm | Fluidigm | 3149001B | 1/100 |
| CD45RA | HI100 | 143 Nd | Fluidigm | 3143006B | 1/100 |
| CCR7 | G043H7 | 159 Tb | Fluidigm | 3159003A | 1/100 |
| *lymphocyte phenotyping* | | | | | |
| CD69 | FN50 | 144 Nd | Fluidigm | 3144018B | 1/100 |
| TBET | 4B10 | 161 Dy | Fluidigm | 3161014B | 1/100 |
| CD28 | CD28.2 | 160 Gd | Fluidigm | 3160003B | 1/100 |
| 4-1BB* | 4B4-1 | 164 Dy | Biolegend | 309802 | 1/50 |
| OX40* | ACT35 | 174 Yb | Biolegend | 350015 | 1/50 |
| CD27 | L128 | 167 Er | Fluidigm | 3167006B | 1/100 |
| ICOS | C398.4A | 148 Nd | Fluidigm | 3148019B | 1/100 |
| CTLA-4 | 14D3 | 170 Er | Fluidigm | 3170005B | 1/100 |
| PD-1 | EH12.2H7 | 155 Gd | Fluidigm | 3155009B | 1/100 |
| CD57 | HCD57 | 163 Dy | Fluidigm | 3163022B | 1/100 |
| Lag3* | 11C3C65 | 151 Eu | Biolegend | 369302 | 1/50 |
| Tim3 | F38-2E2 | 153 Eu | Fluidigm | 3153008B | 1/100 |
| *other phenotyping markers* | | | | | |
| Sox2* | 14A6A34 | 150Nd | Biolegend | 656102 | 1/100 |
| PDL1 | 29E.2A3 | 156 Gd | Fluidigm | 3156026B | 1/100 |
| Ki67 | B56 | 168 Er | Fluidigm | 3168007B | 1/100 |
| cl-caspase3 | 5A1E | 172 Yb | Fluidigm | 3172023A | 1/100 |
| pERK | D13.14.4E | 171 Yb | Fluidigm | 3171010A | 1/100 |
| pAKT | D9E | 152 Sm | Fluidigm | 3152005A | 1/100 |
| pSTAT3 | 4/P-STAT3 | 158 Gd | Fluidigm | 3158005A | 1/100 |
| pSTAT5 | 47 | 147 Sm | Fluidigm | 3147012A | 1/100 |
| pS6 | N7-548 | 175 Lu | Fluidigm | 3175009A | 1/100 |
| Bcl-xl* | E18 | 140 Ce | Abcam | ab199099 | 1/100 |

| | | | | | |
|---|---|---|---|---|---|
| Mcl-1* | Y37 | 154 Sm | Abcam | ab199099 | 1/100 |
| Bcl-2* | clone100 | 157 Gd | WEHI | n/a | 1/100 |

**Supplementary Table 2:** Antibody details in CyTOF antibody panel against human antigens. *indicates in-house conjugated antibodies, prepared at a concentration of 0.1 – 0.4 mg/mL.

| ***human flow cytometry antibodies*** | | | | | |
|---|---|---|---|---|---|
| **antigen** | **clone** | **conjugate** | **manufacturer** | **catalog #** | **dilution** |
| ICOS | C398.4A | BB515 | BD | 565881 | 1/100 |
| CD45 | H130 | AlexaFluor 532 | eBioscience | 58-0459-42 | 1/100 |
| CD45 | HI30 | BV510 | BD | 563204 | 1/100 |
| PD1 | EH12.2H7 | PE | Biolegend | 329906 | 1/100 |
| CD103 | BERact8 | PeDazzle594 | biolegend | 350224 | 1/200 |
| CD4 | OKT4 | Pe-Cy5 | Biolegend | 317412 | 1/50 |
| CD4 | OKT4 | PerCP-Cy5.5 | Biolegend | 317428 | 1/50 |
| CD8 | SK1 | PerCP Cy5.5 | Biolegend | 344710 | 1/50 |
| CD8 | SK1 | Pacific Blue | Biolegend | 344718 | 1/100 |
| HLA-A,B,C | W6/32 | PerCP-eFluor 710 | eBioscience | 46-9983-42 | 1/100 |
| CTLA4 | 14D3 | PE-Cy7 | eBioscience | 25-1529-42 | 1/200 |
| 41BB | 4B4-1 | APC | Biolegend | 309810 | 1/50 |
| CD69 | FN50 | APC-R700 | BD | 565155 | 1/100 |
| EpCAM | 9C4 | Alexa700 | Biolegend | 324244 | 1/50 |
| EpCAM | VU-1D9 | FITC | Stem Cell Technologies | 10109 | 1/40 |
| CD45RA | HI100 | APC-CY7 | Biolegend | 304128 | 1/100 |
| OX40 | BER-ACT35 | BV480 | BD | 746649 | 1/50 |
| CD3 | UCHT1 | BV570 | Biolegend | 300436 | 1/50 |
| CD25 | BC96 | BV605 | Biolegend | 302632 | 1/50 |
| CCR7 | G043H7 | BV650 | Biolegend | 353234 | 1/50 |
| CD27 | L128 | BV750 | BD | 747310 | 1/200 |
| HLA-DR | L243 | BV785 | Biolegend | 307642 | 1/100 |
| CD235a | GA-R2 | PE | BD | 555570 | 1/120 |
| CD140b | 28D4 | PE | BD | 558821 | 1/80 |
| CD31 | WM59 | PE | BD | 555446 | 1/40 |
| ***intracellular antigens*** | | | | | |
| CTLA4 | 14D3 | PE-Cy7 | eBioscience | 25-1529-42 | 1/200 |
| Foxp3 | 206D | BV421 | Biolegend | 320124 | 1/50 |
| Ki67 | B56 | V450 | BD | 561281 | 1/100 |
| Ki67 | B56 | BV786 | BD | 563756 | 1/100 |
| GranzymeB | GB11 | BV510 | BD | 563388 | 1/50 |
| Perforin | dG9 | BV711 | Biolegend | 308130 | 1/200 |

**Supplementary Table 3:** Human antibodies used for flow cytometry analyses.

| ***Nomenclature of T cell populations*** | |
|---|---|
| **population** | **definition** |
| $T_{EM}$ | $CD45^{+}CD3^{+}CD45RO^{+}CD45RA^{-}CCR7^{-}$ |
| $T_{RM}$ | $CD45^{+}CD3^{+}CD69^{+}CD45RO^{+}CD45RA^{-}CCR7^{-}$ |
| $CD4^{+}$ $T_{RM}$ | $CD45^{+}CD3^{+}CD69^{+}CD45RO^{+}CD45RA^{-}CCR7^{-}CD4^{+}CD127^{+}CD25^{lo}$ |
| resident $T_{reg}$ | $CD45^{+}CD3^{+}CD69^{+}CD45RO^{+}CD45RA^{-}CCR7^{-}CD4^{+}CD127^{-}CD25^{hi}$ |
| $CD8^{+}$ $T_{RM}$ | $CD45^{+}CD3^{+}CD69^{+}CD45RO^{+}CD45RA^{-}CCR7^{-}CD8^{+}$ |

**Supplementary Table 4:** Definition of $T_{EM}$ and $T_{RM}$ cells in non-malignant lung tissue. Within tumours, the same gating strategy was used yet cells were termed CD69+ $T_{EM}$ instead of $T_{RM}$.

| ***murine flow cytometry antibodies*** | | | | | |
|---|---|---|---|---|---|
| **antigen** | **clone** | **conjugate** | **manufacturer** | **catalog #** | **dilution** |
| CD45 | 30F-11 | BV570 | Biolegend | 103135 | 1/400 |
| CD3 | 17A2 | AlexaFluor 532 | eBioscience | 58-0032-82 | 1/200 |
| CD4 | RM4-5 | PerCP-Cy5.5 | Biolegend | 100540 | 1/400 |
| CD8a | 53-6.7 | BV650 | Biolegend | 100742 | 1/400 |
| CD44 | IM7 | Alexa700 | Biolegend | 103026 | 1/200 |
| CD62L | MEL-14 | APC-Cy7 | Biolegend | 104428 | 1/200 |
| CD25 | PC61 | BV510 | Biolegend | 102042 | 1/200 |
| ICOS | 7E.17G9 | PerCP-eFluor710 | eBioscience | 46-9942-82 | 1/200 |
| PD1 | 29F1A12 | BV605 | Biolegend | 135219 | 1/100 |
| MHC Class II | M5/114 | PE | WEHI | - | 1/200 |
| CD19 | 6D5 | APC | Biolegend | 115511 | 1/400 |
| CXRC5 | SPRCL5 | Pe-Cy7 | eBioscience | 25-7185-82 | 1/300 |
| Foxp3 | MF-14 | BV421 | Biolegend | 126419 | 1/100 |
| Granzyme B | QA16A02 | PE/dazzle594 | Biolegend | 372215 | 1/100 |
| cl-caspase 3 | C92-605 | V450 | BD | 560627 | 1/100 |
| Ki67 | B56 | BV786 | BD | 563756 | 1/200 |

**Supplementary Table 5:** Murine antibodies used for flow cytometry.

| *human immunotherapeutics* | | | | | |
|---|---|---|---|---|---|
| **antibody** | **clone** | **manufacturer** | **catalog #** | **dosage** | **route** |
| anti-CD3 | OKT3 | WEHI | - | 5 µg/mL | *in vitro* |
| anti-CD28 | CD28.2 | BD | 555725 | 1 µg/mL | *in vitro* |
| human IgG4 | QA16A15 | Biolegend | 403701 | 10-20 µg/mL | *in vitro* |
| hamster IgG | HTK888 | Biolegend | 400966 | 10 µg/mL | *in vitro* |
| nivolumab | - | Selleck Chem | A2002 | 10 µg/mL | *in vitro* |
| ipilimumab | - | Selleck Chem | A2001 | 10 µg/mL | *in vitro* |
| anti-ICOS | C398.4A | Biolegend | 313511 | 10 µg/mL | *in vitro* |
| ***mouse immunotherapeutics*** | | | | | |
| rat IgG2a | 2A3 | BioXCell | BE0089 | 150 µg | i.p. |
| anti-PD1 | RMP1-14 | BioXCell | BE0146 | 150 µg | i.p. |
| anti-ICOS | C398-4A | Biolegend | 313541 | 150 µg | i.p. |
| hamster IgG | HTK888 | Biolegend | 400966 | 150 µg | i.p. |

**Supplementary Table 6:** Immunotherapy antibodies used for human *in vitro* and mouse *in vivo* assays.

| ***human immunohistochemistry antibodies*** | | | | |
|---|---|---|---|---|
| **antigen** | **clone** | **manufacturer** | **catalog #** | **dilution** |
| CD45 | H130 | Biolegend | 304002 | 1/100 |
| ***murine immunohistochemistry antibodies*** | | | | |
| CD8a | polyclonal | SySy | HS-361003 | 1/1000 |
| Ki67 | D3B5 | CST | 12202 | 1/400 |

**Supplementary Table 7:** Antibodies used to detect antigens in histological analyses.

| *datasets* | | | | | | |
|---|---|---|---|---|---|---|
| **name** | **disease type** | **multi-site?** | **# ES patients** | **# NS patients** | **Technology** | **Ref** |
| TCGA | LUAD | no | 420 | 72 | WES<br>RNAseq | Campell 2016<br>Nat Genet |
| TRACERx | LUAD<br>LUSC<br>NSCLC-NOS | yes | 88 | 12 | WES<br>RNAseq | Jamal-Hanjani<br>2017 NEJM |
| LxG | LUAD<br>LUSC<br>SCLC | yes | 16 | 3 | WGS | Leong 2019<br>Oncogene |
| Chen | AIS<br>MIA<br>LUAD | no | 54 | 143 | WES<br>RNAseq | Chen 2019<br>Nat Comms |

**Supplementary Table 8:** Sequencing datasets analysed in this study. LUAD, lung adenocarcinoma; LUSC, lung squamous cell carcinoma; NSCLC-NOS, non-small cell lung cancer not otherwise specified; SCLC, small cell lung cancer; AIS, adenocarcinoma in situ; MIA, minimally invasive adenocarcinoma; WES, whole exome sequencing; RNAseq, RNA sequencing; WGS, whole genome sequencing.
